# Monitoring of drought stress and transpiration rate using proximal thermal and hyperspectral imaging in an indoor automated plant phenotyping platform

**DOI:** 10.1101/2023.08.01.551261

**Authors:** Stien Mertens, Lennart Verbraeken, Heike Sprenger, Sam De Meyer, Kirin Demuynck, Bernard Cannoot, Julie Merchie, Jolien De Block, Jonathan Vogel, Wesley Bruce, Hilde Nelissen, Steven Maere, Dirk Inzé, Nathalie Wuyts

## Abstract

**Background:** Thermography is a popular tool to assess plant water use behavior, as plant temperature is influenced by transpiration rate, and is commonly used in field experiments to detect drought stress. Its application in indoor automated phenotyping platforms is still limited and mainly focuses on differences in plant temperature between genotypes or treatments, instead of estimating stomatal conductance or transpiration rate. In this study, the transferability of commonly used thermography analysis protocols from the field to greenhouse phenotyping platforms was evaluated. In addition, the added value of combining thermal infrared (TIR) with hyperspectral imaging to monitor drought effects on plant transpiration rate (E) was evaluated.

**Results:** The sensitivity of commonly used TIR indices to detect drought-induced and genotypic differences in water status was investigated in eight maize inbred lines in the automated phenotyping platform PHENOVISION. Indices that normalized plant temperature for vapor pressure deficit and/or air temperature at the time of imaging were most sensitive to drought and could detect genotypic difference in the plants’ water use behavior. However, these indices were not strongly correlated to stomatal conductance and E. The canopy temperature depression index, the crop water stress index and the simplified stomatal conductance index were more suitable to monitor these traits, and were consequently used to develop empirical E prediction models by combining them with hyperspectral indices and/or environmental variables. Different modeling strategies were evaluated including single index-based, machine learning and mechanistic models. Model comparison showed that combining multiple thermal infrared indices in a random forest model can improve E prediction accuracy, and that the contribution of the hyperspectral data is limited when multiple indices are used. However, the empirical models trained on one genotype were not transferable to all eight inbred lines.

**Conclusion:** Overall, this study demonstrates that existing TIR indices can be used to monitor drought stress and develop E prediction models in an indoor setup, as long as the indices normalize plant temperature for ambient air temperature or relative humidity.

## Background

Thermography is a popular tool to monitor changes in plant water-use behavior. It is based on the principle that the amount of emitted thermal infrared radiation of a plant relates to its temperature, which in itself depends on incoming radiation and transpiration rate because of the latent heat of evaporation at the leaf surface (1). Transpiration is important in regulating plant temperature for enzymatic processes and in providing nutrients from the soil via the xylem. It is driven by the evaporative demand from the atmosphere and controlled by stomatal conductance (g_s_). Under water-limiting conditions, plants will reduce their E and water loss by closing their stomata resulting in an increase in plant temperature (T_p_). Consequently, the difference between plant and air temperature (T_a_) will be less negative in plants exposed to drought as compared to well-watered plants allowing the detection of this stress.

Thermal infrared sensors have been used in a wide range of studies on abiotic and biotic stresses, such as drought, salinity, and pathogen infection (2–8). The performance in detecting stress strongly depends on stress-induced changes in g_s_ and E. In the case of drought stress, thermography has been implemented in precision agriculture for automating irrigation (9,10). In recent years, thermography also has become a promising tool for plant phenotyping as T_p_ correlates with both water-use behavior and photosynthetic traits, such as E, g_s_, relative water content, water potential (ψ) and non-photochemical quenching (11–15). Thermal infrared sensors have been used to screen for stress tolerant genotypes (4,16) and to monitor the dynamic stomatal responses to abiotic stress (6,17).

Thermography can be applied on different spatial scales from organ to field level (3,4,17,18). Nevertheless, most phenotyping studies have focused on field conditions, while the usability of thermography in indoor phenotyping platforms was much less investigated. The latter allows for the comparison of treatments or genotypes under controlled environmental conditions, which can improve the accuracy of thermal imaging methods, as these are strongly affected by light intensity, temperature, vapor pressure deficit (VPD) and wind speed. Most studies that have implemented thermal infrared imaging in indoor phenotyping platforms have done direct comparisons of T_p_ or have normalized plant temperature by calculating thermal infrared (TIR) indices (6,18–21).

The simplest T_p_ normalization is the Canopy Temperature Depression (CTD) index, which subtracts T_a_ from T_p_ to correct for ambient temperature conditions. This index has been applied to plant disease, heat tolerance and drought stress studies of crops or transgenic plants (7,22,23), as well as for automated irrigation scheduling of wheat in the form of Stress Degree Days (SDD, (24)). An alternative approach to standardize plant temperature is by normalizing it against the temperature of a non-transpiring and fully transpiring leaf or plant. This approach was implemented in the popular Crop Water Stress Index (CWSI) in which CTD was normalized against its maximum (non-transpiring) and minimum (fully transpiring) values. Three methods have been described to calculate the CSWI: the analytical (or energy balance), empirical (or baseline) and direct approach (25). The latter two are more popular, as they require less meteorological information. Instead, the CTD of non- and fully transpiring leaves are estimated with an empirical baseline, a temperature histogram, or with wet and dry reference leaf temperatures (T_wet_ and T_dry_, respectively, (26)). The CWSI has been correlated with leaf water potential, g_s_ and evapotranspiration (8,11,12,18,22,27,28), but in the direct approach it is not linearly related to g_s_ (25). Consequently, the CWSI has mainly been applied to detect stress and not to estimate g_s_ or E. On the other hand, by using the energy balance model, which describes the energy exchange between leaf and environment (1), E and g_s_ can be estimated. This mechanistic approach simplifies and improves the biological interpretation of the relationship between T_p_ and g_s_ or E, but it is based on many meteorological and physical variables and increases the complexity of data processing (17). Simplifications of the energy balance models have been developed, which reduce the amount of required data by incorporating T_wet_ and/or T_dry_ (29). Nevertheless, these adjustments were not sufficient to make the energy balance a widely applied method in phenotyping studies, especially not in indoor phenotyping platforms.

In phenomics, thermal infrared imaging is often combined with fluorescence and hyperspectral imaging, as these systems can capture complementary information on photosynthetic efficiency, leaf anatomy, pigment and water content. The combination of imaging systems is mainly used to monitor multiple physiological traits simultaneously, even though the integration of different data types may produce more robust trait prediction models. The combination of multi/hyperspectral, thermal infrared and/or fluorescence data has been investigated in disease, drought and yield detection in the field (14,19,30–36). Studies that investigated the added value of combining thermal infrared and hyperspectral indices observed an increased prediction or classification accuracy when the different data types were combined in stepwise multiple linear regression, Partial Least Squares regression (PLSR) or Support Vector Machine Classification models (14,32,33).

In this study the transferability of commonly used TIR drought detection and E prediction methods from the field to indoor phenotyping platforms was investigated in a maize drought stress experiment using PHENOVISION, a high-throughput phenotyping platform in semi-controlled greenhouse conditions (37,38). The drought sensitivity of existing and new TIR indices was compared and indices were combined with environmental information to develop E prediction models. In addition, the advantages of combining hyperspectral, thermal infrared and environmental data in physiological trait prediction models were evaluated, as the performance of this approach has only been investigated in field applications (14,32,33). This question was tackled by developing and comparing empirical E prediction models created using different data combinations and modeling approaches, such as linear, lasso, PLSR and Random Forest (RF) regressions. Empirical models are often setup specific, limiting the transferability to other genotypes, species and imaging setups/platforms. To evaluate setup specificity, the empirical models trained on one genotype were applied on seven additional maize inbred lines. The best performing empirical model was also compared to a simplified version of the energy balance approach, which is a setup independent mechanistic model, to determine the most effective approach to estimate E (Additional file: file S1).

## Methods

### Thermal imaging setup and environmental monitoring in the PHENOVISION plant phenotyping system

PHENOVISION is a plant phenotyping platform that automatically irrigates and images 392 plants (built by SMO, Eeklo, Belgium,(39)). Three imaging systems are available: a multi-view red-green-blue (RGB) imaging system, a top view TIR camera and a hyperspectral imaging system (37,38). The TIR camera is positioned in an enclosed imaging cabin to eliminate light from outside. The thermal images are captured using a FLIR SC645 with a 24.6 mm lens (25° × 18.8° FOV) (FLIR Systems Inc., Belgium), which has an accuracy of 2% for objects with a temperature range of −20 to 150 °C. The camera has a fixed top-view (close to nadir) position and is mounted 3.5 - 2.5 m above the plants. At a distance of 3.5 m, the pixels in the 640 X 480 array have a spatial resolution of 5.76 mm^2^. The wavelength band of this camera ranges from 7.5-13 µm. Inside the imaging cabin, light racks containing light emitting diode (LED) lamps are attached to the ceiling and to one wall to provide consistent illumination. The LED lamps provided a photosynthetic active radiation (PAR) light intensity of 74 µmol photons m^−2^s^−1^ and emitted no radiant heat. A Lambertian surface made of aluminum foil and a black metal reference plate are positioned within the field of view of the camera. The aluminum foil is used to measure the reflected temperature, which is the energy emitted by the surroundings of the plant and reflected by the plant. It is required to correct the measured thermal energy of the plant and consequently plant temperature. The black reference plate is connected to a thermocouple to monitor deviations in the camera accuracy over time (camera ‘drift’). In this study, the imaged temperature of the black plate was also used as a surrogate for the temperature of a non-transpiring plant (T_dry_, (3,26)). Thermal data were combined with environmental measurements collected at three different locations in the greenhouse, namely in the growth zone (gz), and inside (in) and outside (out) the cabin (Fig. 1).

**Figure 1:**
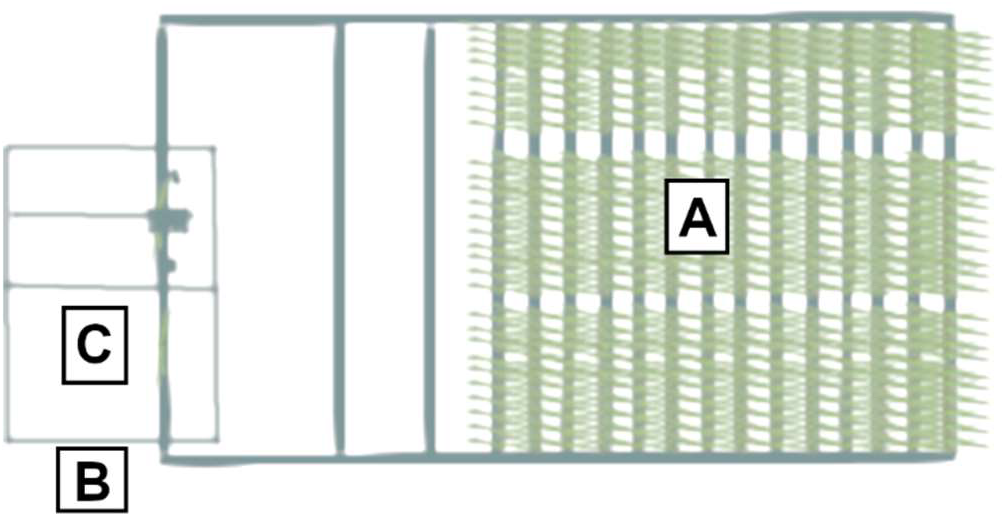
Location of environmental monitoring positions in PHENOVISION. **A** indicates the growth zone. **B** shows the monitoring position outside the imaging cabin. This is the location where the plants are waiting before imaging. **C** indicates the position inside the TIR imaging cabin.

### Experimental setup

Plants were grown in a semi-controlled environment with adjustable T_a_ and relative humidity (RH). During seedling establishment, T_a_ was set at 23-22 °C and vapor pressure deficit was maintained at 1.5 kPa by means of RH adjustments, while a diurnal gradient in environmental conditions was established during plant development (from the V4 - four fully developed leaf stage onwards). This was accomplished by gradually increasing T_a_ from 22 °C at night to 28 °C in the afternoon, which resulted in VPD values ranging from 1.5 to 3 kPa (night/afternoon). A 16/8 h day/night cycle was maintained in the greenhouse using high-pressure sodium vapor lamps, which resulted in an average light intensity (photosynthetically active radiation, PAR) of 280 µmol m^−2^s^−1^. Environmental conditions, including PAR, RH, T_a_, VPD and black sphere temperature (T_BS_), were measured in the growth zone by four weather stations containing a SKH 2053 humidity and temperature sensor, a PAR SKL 2625 sensor (Skye Instruments, UK) and a metal black sphere (Testo BE, Belgium) with a MWTC/MWTC-D thermocouple inside (OMEGA Engineering Ltd., UK). The black sphere temperature represents radiant heat which may be higher than air temperature under influence of solar radiation or heat produced by lamps. Slight variability in PAR and T_a_ was observed within the growth zone which had a negligible effect on the phenotype. Consequently, the gz measurements were averaged. Outside and inside the imaging cabin, RH and T_a_ were monitored by El-USB-2 loggers (Lascar electronics, UK). Maize plants were sown in 7 L plastic pots filled with peat-based soil containing osmocote fertilizer (N. V. Van Israel, Belgium). The plants were fertilized weekly with 40 ml of 200 ppm N Peters Excel CalMag Grower (Everris, The Netherlands) solution.

Two drought stress experiments were performed to determine the effectiveness of TIR imaging for drought detection and E predictions. The first small experiment (DR) investigated the effect of drought on the transpiration rate and stomatal conductance of maize genotype B104, while the second experiment (TF) tested the transferability of thermal image-based phenotyping to other maize genotypes.

During DR, B104 maize plants were positioned randomly on the platform. Four drought treatments with a soil water content of 0.8 (soil water potential < −1500 kPa), 1.0 (< −1500 kPa), 1.4 (± −100 kPa) and 1.6 g H_2_O g^−1^ dry soil (± −40 kPa) and one well-watered treatment with a soil water content of 2.4 g H_2_O g^−1^ dry soil (± −10 kPa) were applied to 104 plants. Twenty plants were randomly assigned to each treatment and were surrounded by border plants, which received a mild water deficit treatment (1.8 g H_2_O g^−1^ dry soil, ± −25 kPa), to remove edge effects. Drought treatment started from germination onwards and lasted until the plants reached V10-11 (end of experiment).

The TF experiment contained a total of 306 plants consisting of eight inbred lines, namely B104, H99, MS71, NC358, OH43, TX303, TZi8 and W153R, which were bordered by 80 B104 plants. This group of inbred lines contained stiff stalk (B104), non-stiff stalk (H99, MS71, OH43 and W153R), tropical-subtropical (NC358 and TZi8) and tropical-subtropical mixed (TX303) genotypes (40) that varied in drought sensitivity, morphology and developmental timing. For each genotype, plants were partitioned between a water-deficit treatment (WD) and a well-watered treatment (WW), which resulted in 19 plants per genotype-treatment combination. All plants were irrigated to a WW soil water content of 2.4 g H_2_O g^−1^ dry soil (soil water potential of −10 kPa) until they reached the V5 (five fully developed leaves) stage after which water was withheld from WD plants until a soil water content of 1.4 g H_2_O g^−1^ dry soil (soil water potential of ± −100 kPa) was attained. Once the WD soil water content was reached, plants were irrigated to sustain this soil water content. During both experiments, RGB, thermal and hyperspectral images were acquired daily, while physiological measurements were collected once per week or at specific sampling time points. The dataset collected during the TF experiment was used to evaluate the drought detectability of thermal TIR indices, to create E prediction models (together with the DR dataset) and to investigate the transferability of indices and models to multiple genotypes.

### Thermal and hyperspectral image processing

Imaging was performed daily between 7.30 AM and 2 PM during DR and TF experiments (14,744 images of eight genotypes). These images were supplemented with 288 afternoon images that were captured on TF sampling days. Thermographic data were processed using the ‘raw2temp’ function of the ‘Thermimage’ R package (41), which converts raw radiation values to temperature using standard equations applied in thermography. This function includes a correction for background long-wave emission, which was estimated using crumpled and flattened aluminum foil (Lambertian surface) with an emissivity of 1. The ‘raw2temp’ function requires the distance between the plant and camera, T_a_, RH and an estimate of leaf emissivity, which was set to 0.96 (42). Deviations in the camera accuracy (camera ‘drift’) were determined per experiment by using a black plate with thermocouple. If a consistent difference higher than 1 °C between the temperature estimated by the camera and the thermocouple was observed, the imaged temperature was adjusted using the median difference in temperature measured by the two devices. After image pre-processing the plant was segmented out of the background by aligning the thermal image with the corresponding segmented RGB image. RGB images were collected using an Allied Vision Technologies Prosilica GE4000C 11-megapixel camera (Allied Vision Technologies GmbH, Germany) equipped with a Canon EF 24 mm f/1.4L II USM lens (Canon Inc., Japan). RGB plant segmentation was performed with a convolutional neural network model developed in python. The segmented and raw RGB images were downscaled from 2672x4008 to 1336x2004 pixel resolution for the thermal image analysis. Both thermal and RGB cameras were previously calibrated using a metal plate with holes or a chessboard (Matlab, Simon Donné, IMEC – IPI – Ghent University), which resulted in intrinsic and extrinsic camera matrices. Due to a slight shift in the RGB camera position the extrinsic matrix could not be utilized. Instead, the translation parameters of the extrinsic matrix were expressed in function of the distance between the RGB camera and the plant, as the height at which the plant was positioned during RGB imaging varied, which affected the alignment of the two images. The transformation matrices were used to align the thermal and RGB images. This required the distance between the plant and the cameras, which was not available for each pixel. Instead, the distance between 2/3 of the plant height and the cameras was used. Using 2/3 of the plant height as an average depth value for all plant pixels did not perform optimally for large plants because the leaves were positioned further apart. To resolve this issue, an erosion was applied on the segmentation filters (binary image) to remove remaining background pixels (‘erode’ function of the ‘mmand’ R package (43)). The median temperature of each plant was calculated and used for drought detection and E predictions.

Hyperspectral data were collected by two pushbroom line-scanners in a second imaging cabin in PHENOVISION. The specifications of this imaging system have been described in Mertens et al. (37). The pre-processing included image calibration, plant segmentation and illumination classification. The illumination classification was not able to remove all the light effects, because the distance between the white reference plate, which is used to calibrate the hyperspectral images, and the top of the plant was too big for large plants. This issue was solved by correcting the reflectance distribution of a plant (pixel level) to the distribution it would have had at the white reference height tile in the PHENOVISION imaging cabin (at 1.2 m from the imaging system). Three plants were therefore imaged at different lift heights and the relationship between average reflectance and the distance between plant and camera was determined (Fig. 2). This relationship was then used to adjust the reflectance distribution on a pixel level with the following formula.

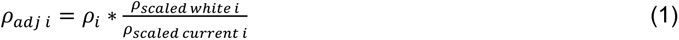

**Figure 2:**
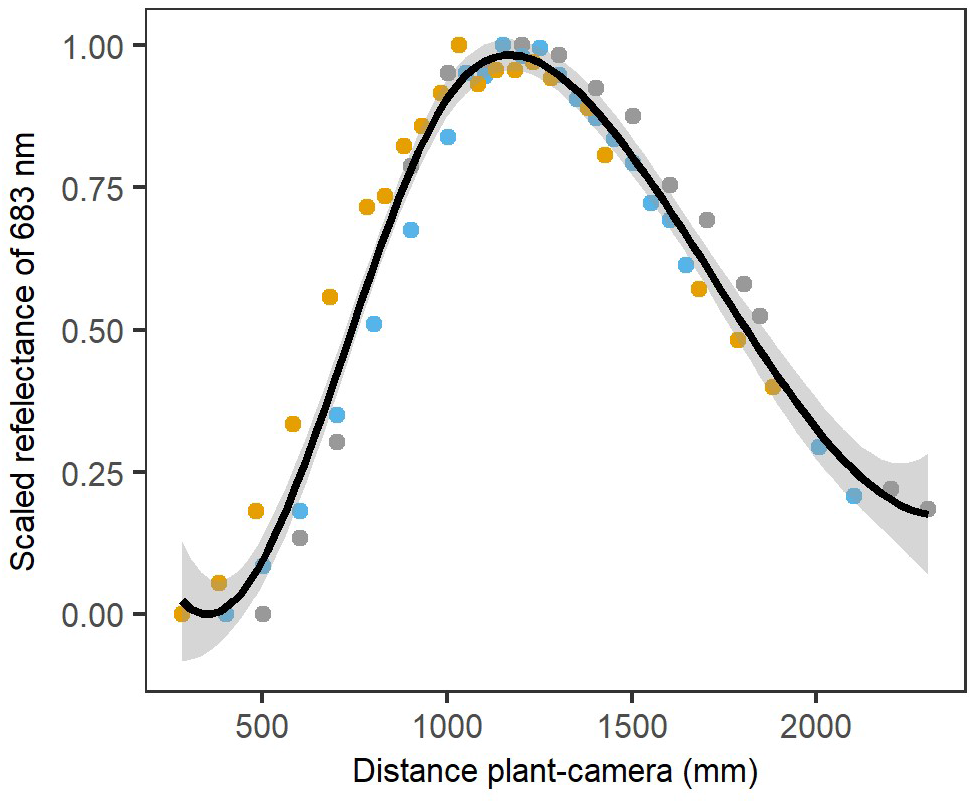
Relationship between reflectance and the distance between the plant and the camera. Average plant reflectance was scaled between zero and one to remove biological variation between the plants. Three plants were used to determine this relationship, indicated with gray, yellow and blue dots. The average relationship and the 95% confidence interval are indicated by a black line and gray shading, respectively.

In this equation, *ρ_adj i_* is the adjusted reflectance of wavelength i, *ρ_i_* corresponds to the unadjusted reflectance, *ρ_scaled white i_* represents the scaled average reflectance of the plant positioned at the white reference and *ρ_scaled current i_* is the scaled average reflectance at the actual distance from the camera. The reflectance distribution correction was performed before illumination classification and reflectance averaging. To simplify the analysis, only one light class (intermediate light class) was selected based on the percentage of pixels per plant it contained during all experiments and the ability to detect drought (37).

### Physiological trait measurement

Gas-exchange measurements were collected on a weekly basis during DR. Five plants per irrigation treatment (total of 25) were measured 5-10 min before imaging, which occurred between 7.30 AM and 2 PM. The gas-exchange measurements were collected using a portable LICOR 6400-XT infrared gas analyzer (LI-COR Biosciences, USA). Within the leaf chamber, a steady state CO_2_ level of 400 µmol mol^−1^ was maintained, while temperature and PAR were adjusted to the greenhouse temperature (25-31 °C) and PAR (50-700 µmol photons m^−2^s^−1^).

The monitoring approach of DR was also applied during the TF experiment, where fluorescence and gas-exchange measurements were collected on three plants per genotype-treatment combination (total of 48 plants). Additional physiological measurements were collected at 10 days after V5 (acute drought phase) and ± two days after V13 (13 fully developed leaves). Nine plants per genotype-treatment combination (144 plants) were used for sampling producing 288 additional physiological measurements, which were collected in the afternoon (1-6 pm). Non-destructive fluorescence and gas-exchange measurements consisted of effective quantum yield of photosystem II (ϕ_ps2_), energy harvesting efficiency by oxidized PSII (F_v_’/F_m_’), E and g_s_ of H_2_O. In addition, leaf ψ of a top leaf, which was clearly visible in the image, was destructively measured 5-10 min after imaging using a PMS model 1000 pressure chamber (PMS Instrument Company, USA). Six plants per genotype-treatment combination were selected to monitor ψ on a weekly basis during the whole TF experiment. Each individual plant was measured every two weeks resulting in 48 measurements per week.

### Indices

Thermal infrared indices were calculated to detect drought and estimate transpiration rate for different maize genotypes. These indices combined T_p_ with T_dry_ or environmental measurements collected in the growth zone, and inside or outside the imaging cabin (Fig. 1). Six existing and two new indices, T_BS_-T_p_ and temperature ratio index (TRI), were evaluated (Table 1). T_BS_-T_p_ is an adaptation of T_dry_-T_p_ in which T_dry_ is replaced by T_BS_. This was tested as both temperature variables (T_dry_ and T_BS_) measured the combined effect of incoming radiation and T_a_ on an object’s temperature. The temperature ratio index (TRI) was evaluated because it could detect drought treatments and correlated with g_s_ and E. The ICWSI and CWSI indices were calculated using the baseline approach. This requires the estimation of the maximum temperature or minimum CTD of a non-transpiring plant and the minimum temperature or maximum CTD of a fully transpiring plant. The maximum temperature was assumed to be T_a_ + 3 °C based on the maximum CTD observed in this study and literature (44), while T_p_ or CTD of a fully transpiring plant was determined with a baseline that related T_p_ or CTD to VPD for each genotype (12 WW plants, 38 ± 4 images per genotype). The baseline models were created using the ‘lm’ function of the ‘stats’ R package (45). The ICWSI used a development corrected baseline, while for the CWSI two adaptations were implemented that corrected the index for developmental stage or non-constant environmental conditions (Additional file: Fig. S1). Therefore, baselines were created for juvenile and mature plants separately (ICSWI, CWSI_dev_) or by adding T_a_ as a predictor to the baseline models (CWSI_Ta_, (25)).

**Table 1:** Thermal infrared and hyperspectral reflectance indices evaluated in this study.

| Index | Formula | Reference |
| --- | --- | --- |
| **Thermal Infrared** |  |  |
| Canopy Temperature Depression (CTD) | $CTD = T_p - T_a$ | (47) |
| Canopy Stress Index (CSI) | $CSI = \frac{T_p - T_a}{VPD}$ | (48) |
| Idso's Crop Water Stress Index (ICWSI) | $ICWSI = \frac{T_p - T_{min}}{T_{max} - T_{min}}$ | (49) |
| Crop Water Stress Index (CWSI) | $CWSI = \frac{(T_p - T_a) - (T_p - T_a)_{max}}{(T_p - T_a)_{min} - (T_p - T_a)_{max}}$ | (44,50) |
| Simplified Stomatal Conductance Index ($T_{dry} - T_p$) | $T_{dry} - T_p$ | (51) |
| Temperature Ratio Index (TRI) | $TRI = \frac{T_p}{T_a}$ | this study |
| $T_{BS} - T_p$ | $T_{BS} - T_p$ | this study |
| **Hyperspectral** |  |  |
| 1<sup>st</sup> derivative Simple Ratio index $_{660/1040}$ (dSR$_{660/1040}$) | $dSR_{660/1040} = \rho'_{660}/\rho'_{1040}$ | (52) |
| Normalized Difference index $_{1425/2145}$ (ND$_{1425/2145}$) | $ND_{1425/2145} = \frac{(\rho_{1425} - \rho_{2145})}{(\rho_{1425} + \rho_{2145})}$ | (53) |
| Water Band index (WBI) | $WBI = \rho_{900}/\rho_{970}$ | (54) |
| Simple ratio index $_{1440/1460}$ (SR$_{1440/1460}$) | $SR_{1440/1460} = \rho_{1440}/\rho_{1460}$ | (53) |
| Normalized Difference Water Index (NDWI) | $NDWI = \frac{\rho_{857} - \rho_{1241}}{\rho_{857} + \rho_{1241}}$ | (55) |
| Relative Moisture Percentage 1483/1430 (RMP$_{1483/1430}$) | $RMP_{1483/1430} = \frac{\rho_{1483}}{\rho_{1430}}$ | (56) |
| Water Content Index (WCI) | $WCI = \frac{(\rho_{686} - \rho_{955})}{(\rho_{955} - \rho_{548})}$ | (37) |
| Water Potential index 2 (WP$_2$) | $WP_2 = \frac{(\rho_{665} + \rho_{1457})}{(\rho_{715} + \rho_{1457})}$ | (37) |
| Ratio index 953/492 (R$_{953/492}$) | $R_{953/492} = \frac{\rho_{953}}{\rho_{492}}$ | (37) |
| Normalized Difference index 1407/1862 (NDI$_{1407/1862}$) | $NDI_{1407/1862} = \frac{\rho_{1407} - \rho_{1862}}{\rho_{1407} + \rho_{1862}}$ | (37) |

*Reflectance wavelengths are indicated by $\rho$ .*

Hyperspectral indices were evaluated by determining their relationship with E and g_s_. Ten indices were selected for this analysis of which six have been correlated with E and four with water content or ψ (Table 1). Correlations of indices and environmental data with E and g_s_ were calculated using the ‘cor.mtest’ function of the ‘corrplot’ R package (46). Indices and environmental data that showed strong correlations were subsequently used to create linear E prediction models.

### Transpiration rate modelling

Data on T_p_ and indices, and environmental and hyperspectral data were used to predict E. Different combinations of datasets and modeling approaches were evaluated. Linear models were created to relate T_p_, indices and environmental data with E. Index-based linear models had E as dependent variable and an index as independent variable, while correlation-based linear models combined one strongly correlating index with environmental data (‘lm’ function of ‘stats’ R package). The significant contribution of different predictors to the linear model was evaluated using the Chi-square test of the ‘anova’ function (‘stats’ R package). To investigate if combining multiple indices with T_p_ and environmental data could improve prediction accuracy, RF, lasso and stepwise selection models were created using the ‘randomForest’, ‘glmnet’ and ‘olsrr’ R packages, respectively (57–59). The RF model hyperparameters were fine-tuned by leave-one-out cross validation (LOOCV) and the optimal number of predictors was determined with the ‘VSURF’ R package (60). To combine thermal, environmental and hyperspectral data, correlation-based linear models that combine TIR and hyperspectral indices, RF, lasso, stepwise selection and PLSR models were evaluated (61). The PLSR models were created with the PLSR function of the ‘pls’ R package as described in Mertens et al. (37). The E prediction models were developed using ordered quantile normalized data (‘orderNorm’ function, ‘bestNormalize’ R package, (62)). The prediction accuracy of the models was evaluated by calculating the ‘out-of-bag’ test Mean Absolute Percent Error (MAPE), the Root Mean Square Error (RMSE) and R-squared (R^2^) (‘postResample’ function, ‘caret’ R package, (63)) of 100 bootstrap samples. A final model was created using the whole B104 dataset. This model was subsequently used to test the transferability of the models to different genotypes.

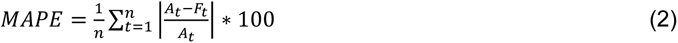

The description and results of the mechanistic energy balance model that was evaluated during this study can be found in the additional file: file S1.

### Drought detection and statistics

Drought detectability of TIR indices was evaluated by comparing when significant differences between the WW and WD treatments were first detected and for how many days. To take variation in the start of the drought into account, time was expressed as days after V5 (start drought). Treatment differences were tested for each day and for each genotype (start experiment: n=19, end: n=5) with a factorial ANOVA using the ‘lsmeans’ R package (64). Factorial ANOVA was also used to test treatment differences of E, g_s_, ψ and ϕ_ps2_ for every group of four days and genotype (n=3, can be higher on sampling days). The days were grouped together because measurement cycles were spread over a few consecutive days. The data were split into morning (<1 PM) and afternoon (>1 PM) measurements to take the time-of-day effect on physiology into account. The TIR index data were not split into morning and afternoon, as the majority of the images were collected in the morning. All the p-values were corrected using the sidak method (‘MHTmult’ R package, (65)).

Genotypic differences in the sensitivity of TIR indices and physiological measurements to drought stress effects were tested for each day after V5. The ‘lsmeans’ R package was used to test genotypic differences in treatment contrasts with the ‘contrast’ function. The genotype analysis of the physiological traits could only by performed at 10 days after V5 (n=8-10 for each treatment), because the sample size was too small on the other days. The p-values of this analysis were also corrected using the sidak method (‘MHTmult’ R package, (65)).

## Results

### Drought affected the water use behavior of B104 maize

Drought stress effects on plant physiology have been widely studied, as they can have a major impact on plant performance and yield (66). In the TF experiment, drought was applied on B104 maize from the V5 developmental stage onward. The drought treatment consisted of two stages, namely an acute drought stage in which water was withheld from the plant, and a steady state drought, during further development, when the plant was kept at a lower soil water content. Plant physiology was most strongly affected during and shortly after the acute drought stage in which a decrease in the g_s_, E, ϕ_ps2_ and ψ was observed in the WD treatment (Fig. 3A). The treatment differences were only significant for the afternoon ψ and g_s_ measurements collected at 10 days after the onset of drought. The absence of significant differences in the other physiological traits and time points may be related to the limited number of available data points. The difference between WD and WW plants became less pronounced during the steady-state drought as the WD plants adapted to the lower soil water content. This adaptation included a decrease in biomass, as the WD B104 plants had on average 39% lower fresh weight at silking compared to WW plants (WW: 615.3±70.9 g, WD: 375.3±34.2 g) resulting in lower water requirements.

**Figure 3:**
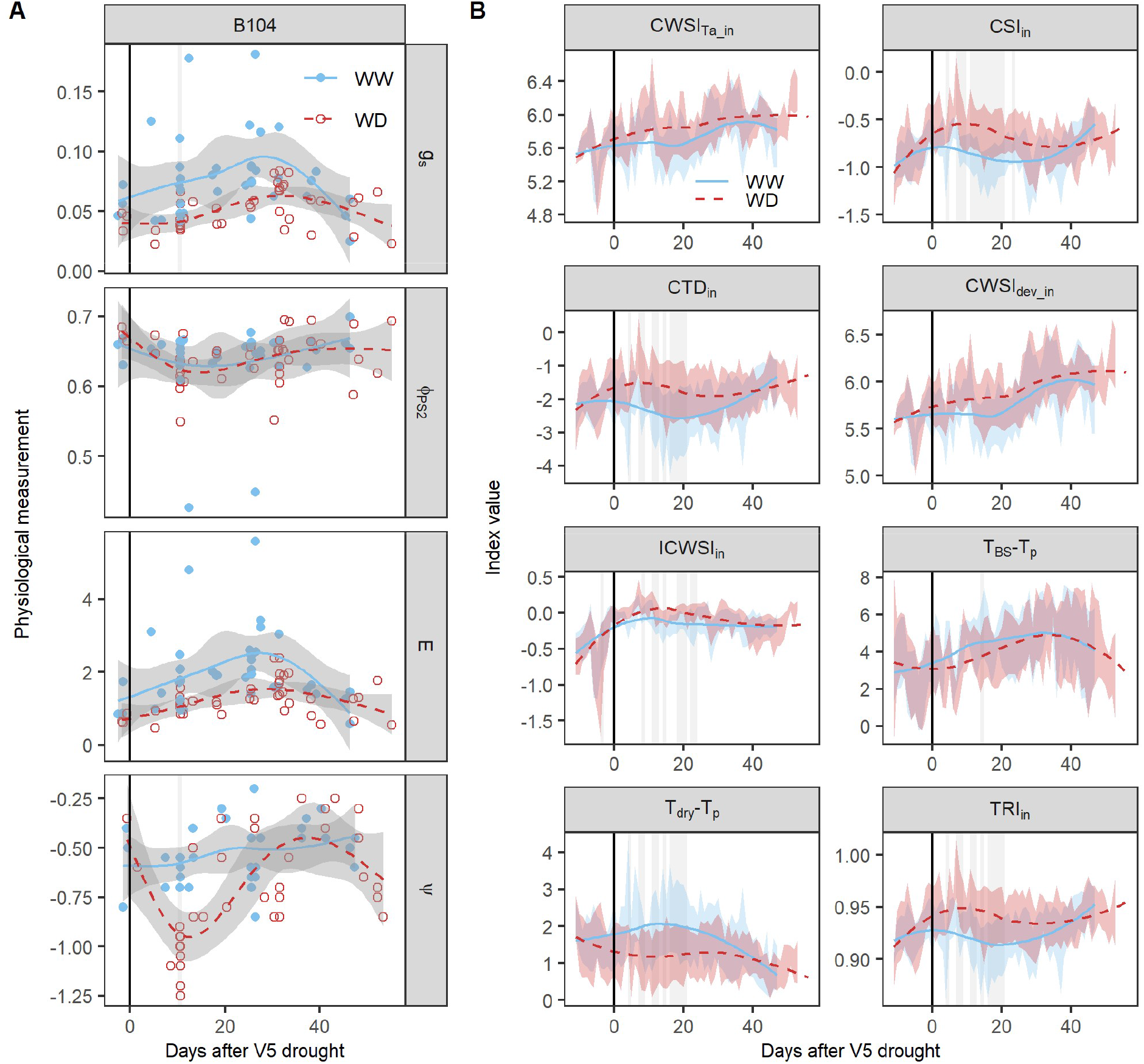
Responses of physiological traits and thermal infrared indices to drought. **A**, drought effects on plant physiology during the TF experiment. The B104 genotype and four physiological traits, stomatal conductance (g_s_, mol H_2_O m^−2^s^−1^), effective quantum yield of photosystem II (ϕ_ps2_), transpiration rate (E, mmol H_2_O m^−2^s^−1^) and leaf water potential (ψ, MPa), were selected for this analysis. **B**, responses of thermal infrared (TIR) indices to drought. B104 maize plants (n_WD_: 19, n_WW_: 19) were imaged daily and TIR indices were calculated using the formulas described in Table 1. Well-watered (WW) and water-deficit (WD) maize plants were monitored from V4 until the silking stage. The average trends of the WW and WD treatments are indicated by blue solid and red dashed lines, respectively. The 95% confidence interval of the average is represented by gray shading (**A**) and the standard deviation by blue and red shading for the WW and WD treatments, respectively (**B**). Individual measurements of the plants are visualized by blue dots (WW) and red circles (WD). The black vertical line indicates the start of the drought treatment. The days on which significant treatment differences were observed are marked with a light gray vertical shading behind the average trend and dots (P<0.05).

### Thermal infrared indices can detect drought in an automated phenotyping platform

All indices, that normalized T_p_ using environmental variables, were calculated for each of the three monitoring positions to determine which one was most suitable for drought detection. The three monitoring positions showed slight differences in VPD and T_a_. During the TF experiment VPD and T_a_ measurements were highest outside the imaging cabin (VPD_out_: 2.2 kPa, T_a_out_: 27 °C), followed by the positions inside the cabin and at the growth zone (VPD_in_: 1.9 kPa, T_a_in_: 25.53 °C, VPD_gz_: 1.8 kPa, T_a_gz_: 25 °C) (Additional file: Fig. S2). Consequently, indices produced slightly different values depending on the monitoring position with those using VPD and/or T_a_ measured inside the imaging cabin performing better (Additional file: Fig. S2, Table 2). The indices that included VPD and/or T_a_ collected outside the cabin or in the growth zone were excluded to simplify the drought detection analysis.

**Table 2:** The effect of environmental measurement location on the drought sensitivity of the thermal infrared indices.

| Index | Number of days with significant treatment differences |  |  |
| --- | --- | --- | --- |
|  | Inside cabin | Outside cabin | Growth zone |
| CSI | 15 | 4 | 1 |
| CTD | 11 | 7 | 1 |
| $T_{dry}-T_p$ | 11 | \ | \ |
| TRI | 11 | 7 | 1 |
| ICWSI | 9 | 1 | 0 |
| CWSI | 0 | 0 | 0 |
| $CWSI_{T_a}$ | 1 | 0 | 0 |
| $T_{BS}-T_p$ | 0 | 0 | 1 |

*The effect of location on drought sensitivity was assessed by means of the number of days that showed significant differences in index values between the well-watered and water-deficit treatment. Missing data are indicated by \, which resulted from the fact that the $T_{dry}-T_p$ index only used measurements collected inside the imaging cabin.*

Thermal indices showed a similar pattern in drought responsiveness as the water-use behavior measurements (E, g_s_, ψ, Fig. 3A), with stronger differences between treatments during the acute drought period and a recovery during the steady-state drought. The CSI showed the strongest drought stress effects with significant treatment differences visible from 4 until 23 days after the onset of drought (Fig. 3B). This was followed by TRI, CTD and T_dry_-T_p_ in which the drought stress effects lasted from 4 to 20 days after the onset of drought, while for ICWSI it lasted from 8 to 23 days. CWSI_dev_, CWSI_Ta_ and T_BS_-T_p_ were less or not sensitive to drought as significant effects were absent (CWSI_Ta_) or only visible on a few days (CWSI_dev_ and T_BS_-T_p_, Fig. 3B). The thermal indices were not only influenced by soil water deficit, but also showed a trend during the morning and early afternoon with lower CSI and TRI values for WW plants around noon compared to the morning (Fig. 4). These diurnal changes corresponded with an increase in VPD and T_a_ in the greenhouse and a decrease in CTD. The decrease was less pronounced for WD plants, which often resulted in larger treatment differences around noon than in the morning. Trends were also visible in the physiological measurements, where a decrease in ψ was observed during the morning, which was more pronounced for WD compared to WW plants (Fig. 4G). Overall, TIR indices were able to detect drought stress effects in an automated phenotyping platform as long as environmental data, such as T_a_ and VPD, were used to normalize T_p_.

**Figure 4:**
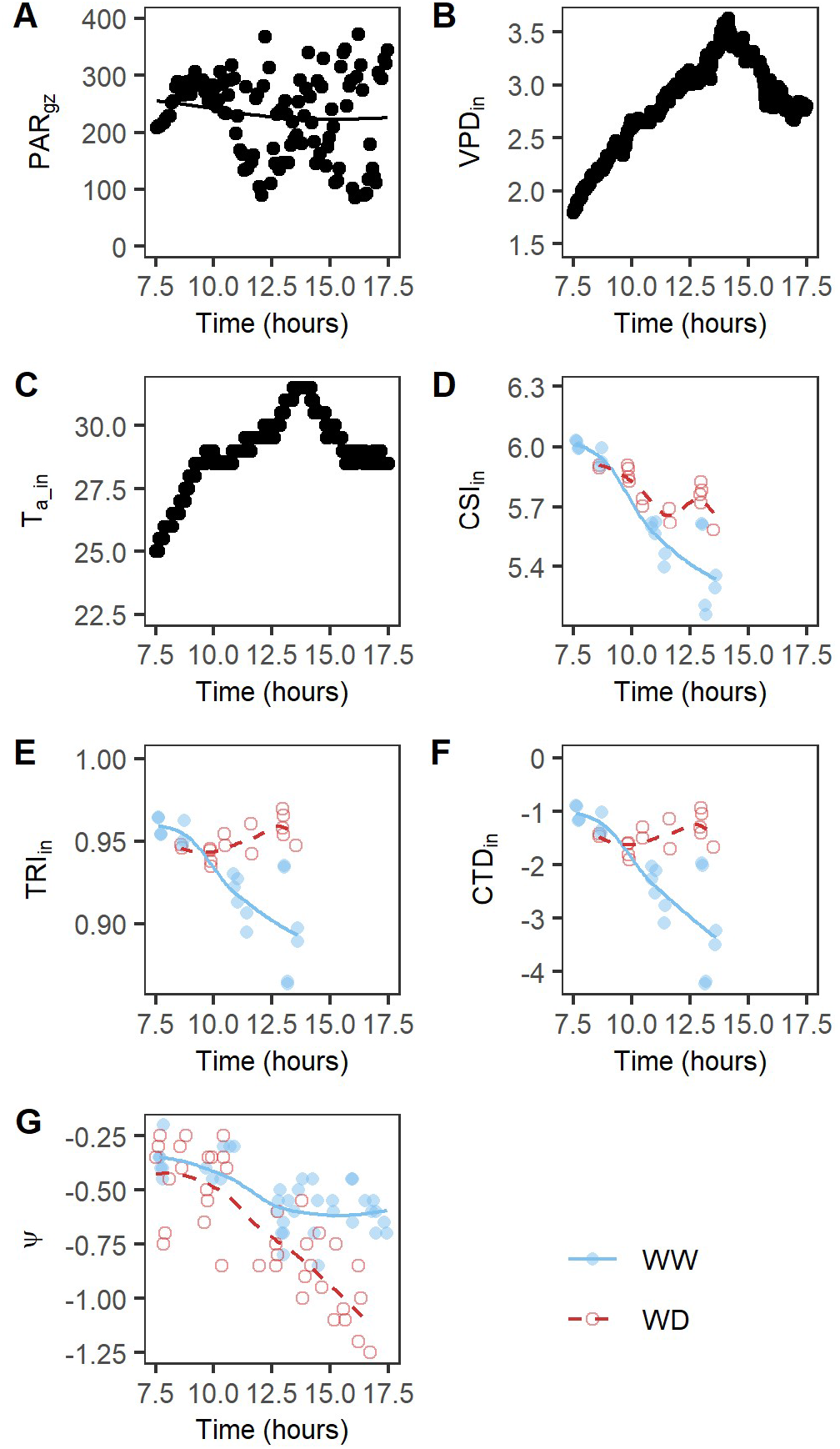
Daytime trend in environmental data, thermal infrared indices and leaf water potential of B104 plants. For the environmental data and indices, one day during the acute drought period was selected that corresponded with ±5 days after the onset of drought. For leaf water potential (ψ), all measurements collected during the experiment were used because of the limited number of B104 measurements per day. Average and individual environmental measurements (A-C) are shown with black lines and black dots, respectively. The average index and ψ values of the well-watered (WW) and water-deficit (WD) treatment are represented by a blue line and a red dashed line, respectively (D-G). The individual plants are visualized with blue dots (WW) and red circles (WD). Daytime patterns of meteorological data: (**A**) PAR in the growth zone (PAR_gz_), (**B**) VPD inside the cabin (VPD_in_), and (**C**) air temperature inside the cabin (T_a_in_). Morning patterns of TIR: (**D**) CSI_in_ index, (**E**) TRI_in_ index, and (**F)** CTD_in_ index. (**G**) Daytime trend in leaf ψ.

### Transpiration rate can be predicted using a combination of environmental data and thermal indices

Besides drought detection, TIR indices can be used to monitor g_s_ and E. In the PHENOVISION setting, relatively strong correlations with g_s_ and E were observed for T_dry_-T_p_ (r_E_=0.61, r_gs_ =0.50, P<0.05), CWSI_Ta_ (r_E_=-0.61, r_gs_=-0.58, P<0.05) and CTD (r_E_=-0.63, r_gs_=-0.64, P<0.05) (Fig. 5B). In the case of CWSI_Ta_ and CTD, the highest correlations were found when environmental data measured outside the imaging cabin were used. This is also the location where the gas exchange measurements were performed. Consequently, also E and g_s_ showed the highest correlations with environmental data collected at this position (Fig. 5A). PAR and T_BS_, which were only measured in the growth zone, were also significantly correlated with E and g_s_ (r_E-PAR_ =0.54, r_E-TBS_= 0.58, r_gs-PAR_ = 0.54,r_gs-TBS_ = 0.54, P<0.05). The correlated environmental data and indices were then combined to create empirical models that can predict E.

**Figure 5:**
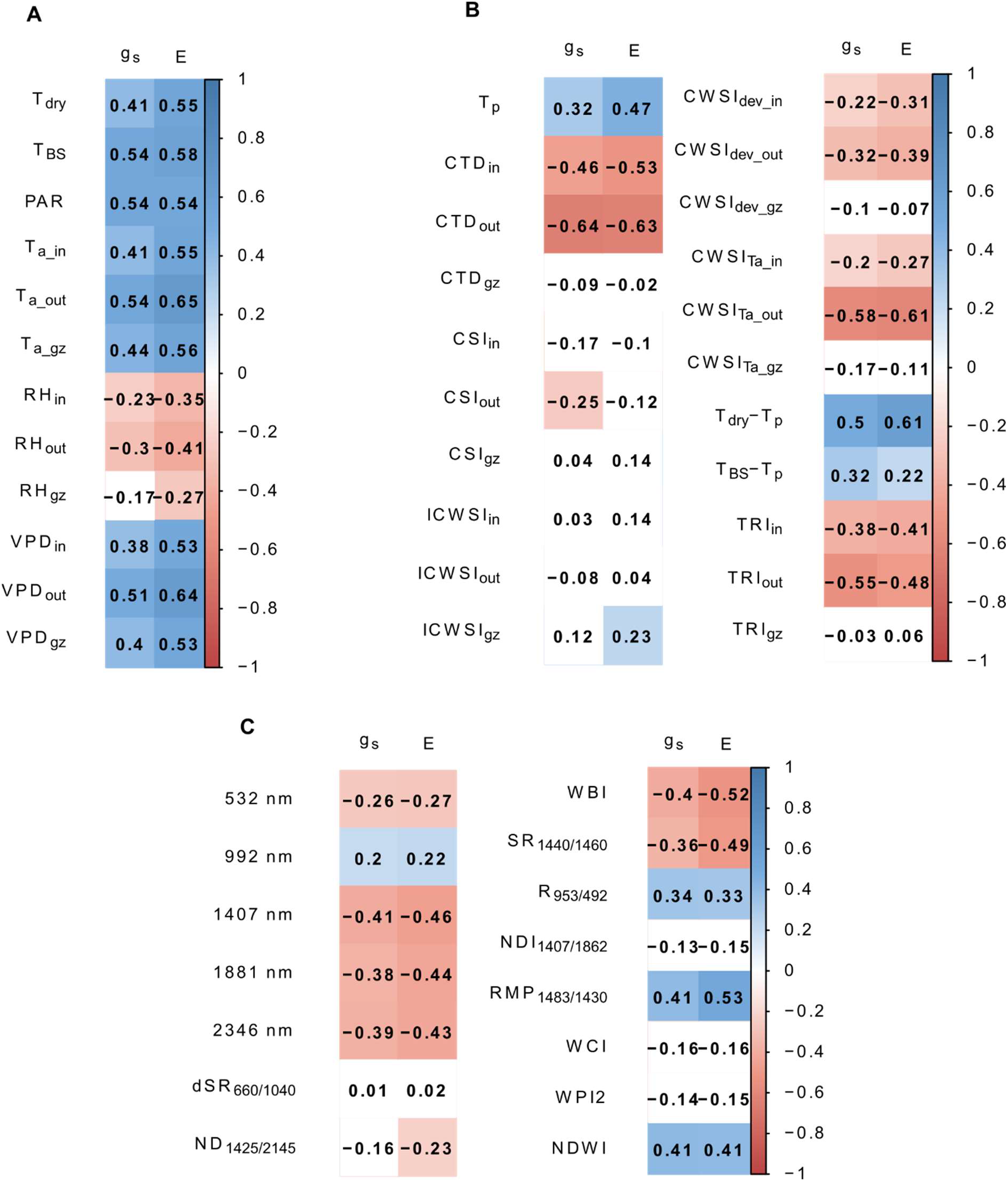
Correlation of stomatal conductance (g_s_) and transpiration rate (E) with measured independent variables. The independent variables included (**A**) environmental variables: temperature of a non-transpiring plant (T_dry_), photosynthetic active radiation (PAR), black sphere temperature (T_BS_), air temperature (T_a_), relative humidity (RH) and vapor pressure deficit (VPD) measured in the growth zone (gz), and inside (in) and outside (out) the cabin, (**B**) thermal infrared indices (Table 1), and (**C**) relative reflectance at 532, 992, 1407 and 1881 nm and hyperspectral indices (Table 1). Significant (P<0.05) correlations are indicated in blue (positive) and red (negative).

The indices with the strongest correlations, CWSI_Ta_out_, CTD_out_ and T_dry_-T_p_ (Fig. 5B and Additional file: Fig. S3), had the highest prediction accuracy of all the index-based models with an RMSE of 0.68, 0.66 and 0.61 and an R^2^ of 0.39, 0.44 and 0.41, respectively (Table 3, Fig. 6). The test prediction accuracy of these models was improved by combining the indices with environmental data, such as VPD_out_, PAR and T_BS_ (RMSE_CWSI_TaOut-env_ = 0.57, RMSE_CTD_Out-env_ = 0.60, RMSE_Tdry-Tp-env_ = 0.56, R^2^_CWSI_TaOut-env_ = 0.52, R^2^_CTD_Out-env_ = 0.57 and R^2^_Tdry-Tp-env_ = 0.55, Table 4, Fig. 6). VPD_out_ was the only environmental variable that improved the prediction accuracy of all three index-based models. PAR and T_BS_ were added to the index-based models because they relate to the incoming radiation, which may additionally increase T_p_ independently of T_a_.

**Figure 6:**
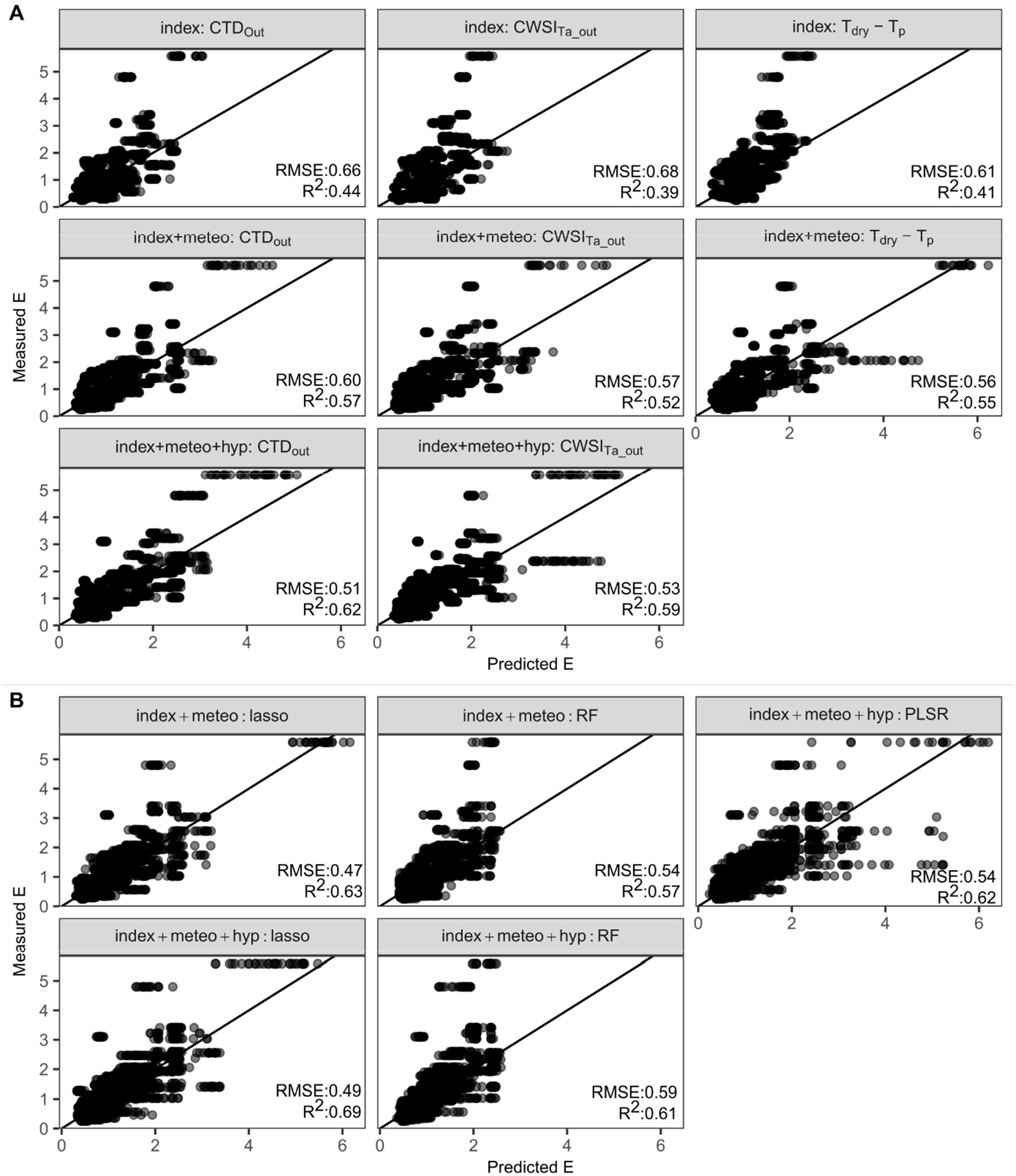
Relationship between measured and predicted transpiration rate (E). E was measured with a portable LICOR infrared gas analyzer. (**A**) The first row visualizes the test prediction accuracy of thermal infrared (TIR) index-based models (CTD_out_, CWSI_Ta_out_, and T_dry_-T_p_), while the second row shows the performance of these indices when they are combined with environmental data not included in the index (Table 4). The third row contains two figures that show the accuracy of models that combine one TIR index (CTD_out_ or CWSI_Ta_out_) with environmental data and hyperspectral indices (RMP_1483/1430_, WBI). This type of model was not created for the T_dry_-T_p_ index as hyperspectral indices did not significantly contribute to this model (Chi-square test, P<0.05). (**B**) Prediction accuracy of the lasso, RF and PLSR models that combine multiple TIR indices with hyperspectral wavelengths and/or environmental data. The semi-transparent black dots represent the predicted and measured E of the 100 bootstrap samples, while the black line shows the one-to-one relationship. The test RMSE and R^2^ is added to each scatterplot.

**Table 3:** The prediction accuracy of normalized thermal infrared indices.

| Index | RMSE | $R^2$ | MAPE (%) |
| --- | --- | --- | --- |
| CTD<sub>in</sub> | 0.66 | 0.32 | 55 |
| CTD<sub>out</sub> | 0.66 | 0.44 | 44 |
| CTD<sub>gz</sub> | 0.8 | 0.02 | 62 |
| CSl<sub>in</sub> | 0.8 | 0.01 | 61 |
| CSl<sub>out</sub> | 0.73 | 0.02 | 62 |
| CSl<sub>gz</sub> | 0.79 | 0.02 | 60 |
| ICWSl<sub>in</sub> | 0.78 | 0.05 | 58 |
| ICWSl<sub>out</sub> | 0.86 | 0.02 | 61 |
| ICWSl<sub>gz</sub> | 0.8 | 0.05 | 57 |
| CWSl<sub>in</sub> | 0.76 | 0.1 | 61 |
| CWSl<sub>out</sub> | 0.77 | 0.13 | 60 |
| CWSl<sub>gz</sub> | 0.85 | 0.03 | 62 |
| CWSl<sub>Ta_in</sub> | 0.8 | 0.11 | 62 |
| CWSl<sub>Ta_out</sub> | 0.68 | 0.39 | 51 |
| CWSl<sub>Ta_gz</sub> | 0.79 | 0.06 | 65 |
| $T_{dry}-T_p$ | 0.61 | 0.41 | 50 |
| $T_{BS}-T_p$ | 0.8 | 0.04 | 57 |
| TRl<sub>in</sub> | 0.72 | 0.17 | 60 |
| TRl<sub>out</sub> | 0.7 | 0.21 | 54 |
| TRl<sub>gz</sub> | 0.76 | 0.03 | 61 |

*Index-based models were created for each index and the prediction accuracy was evaluated by* *calculating the median test RMSE, $R^2$ and MAPE of 100 bootstrap samples.*

**Table 4:** Prediction accuracy of transpiration rate prediction models that use one thermal infrared index.

| RMSE | $R^2$ | MAPE | Selected Variables | Method |
| --- | --- | --- | --- | --- |
| Linear: single TIR index |  |  |  |  |
| 0.56 | 0.55 | 41 | $T_{dry}-T_p, T_{BS}, VPD_{out}, interaction_{T_{dry}-T_p, VPD_{out}}$ | Linear: index + env |
| 0.57 | 0.52 | 39 | $CWSl_{Ta_{out}}, VPD_{out}, PAR_{gz}, T_{BS}$ | Linear: index + env |
| 0.60 | 0.57 | 37 | $CTD_{out}, VPD_{out}, PAR_{gz}$ | Linear: index + env |
| 0.53 | 0.59 | 35 | $CWSl_{Ta_{out}}, T_{BS}, WBI$ | Linear: Index + env +<br>hyperspectral index |
| 0.51 | 0.62 | 33 | $CTD_{out}, PAR_{gz}, RMP_{1483/1430}, WBI$ | Linear: Index + env +<br>hyperspectral index |

To evaluate the impact of combining different indices and environmental data on the prediction accuracy of E, stepwise selection, lasso and RF models were developed and compared with the single index models. This comparison showed that including multiple indices and environmental variables can improve the prediction accuracy even further. The lasso model had the highest accuracy of all TIR models with a median test RMSE=0.47 and test R^2^=0.63 (Table 5, Fig. 6). Interpreting the variables of a lasso model is difficult as it removes redundant variables, which may have biological relevance. Consequently, the variables of the RF model (second-best) were evaluated (Table 5). RF selected four environmental variables (T_a_out_, VPD_gz_, PAR_gz_ and T_BS_gz_) and 11 indices (CTD_out_, CSI_in_, CSI_out_, ICWSI_in_, CWSI_dev_in_, CWSI_dev_out_, CWSI_dev_gz_, CWSI_Ta_gz_, T_BS_-T_p_, TRI_in_ and TRI_gz_). Most of these variables were significantly correlated with E, except for the CSI indices, ICWSI_in_ and the gz indices. The RF model incorporated environmental data of all three monitoring positions directly or indirectly (through indices). This may indicate that all three monitoring positions capture relevant information to predict E in PHENOVISION. The value of the different monitoring positions may be explained by the fact that the plants were not stabilized and acclimatized to the environment of the cabin during TIR imaging. In PHENOVISION maize plants can adjust to the growth zone environment and the waiting area outside the imaging cabin, as they remain at these positions for some time, while in the imaging cabin they are immediately imaged to avoid acclimatization to an irrelevant environment. Consequently, the transportation of the plants around the platform increases the need of monitoring the environment at multiple locations.

**Table 5:**
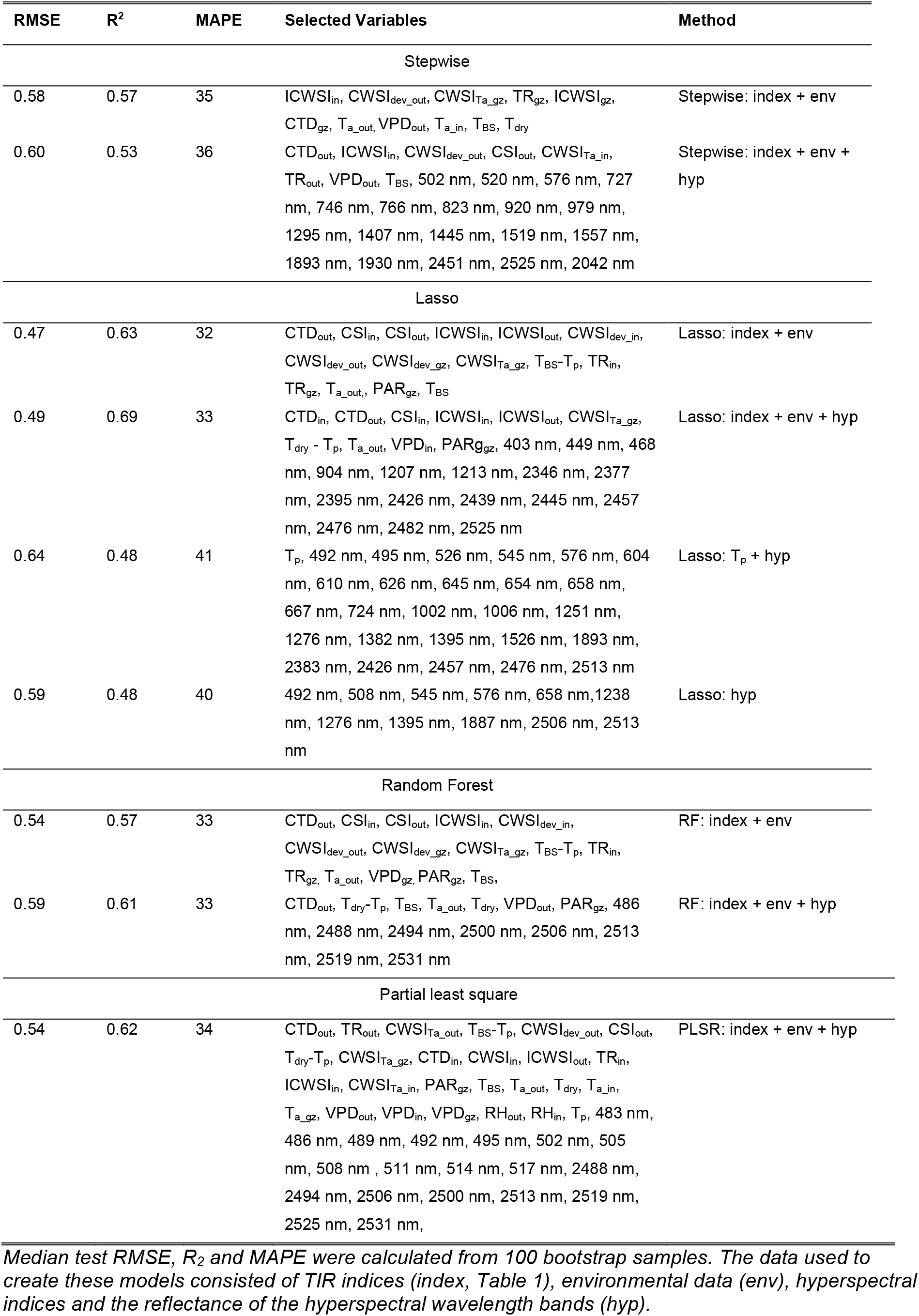
Accuracy of transpiration rate prediction models based on multiple thermal infrared indices.

| RMSE | R<sup>2</sup> | MAPE | Selected Variables | Method |
| --- | --- | --- | --- | --- |
| Stepwise |  |  |  |  |
| 0.58 | 0.57 | 35 | ICWSI<sub>in</sub>, CWSI<sub>dev_out</sub>, CWSI<sub>Ta_gz</sub>, TR<sub>gz</sub>, ICWSI<sub>gz</sub>, CTD<sub>gz</sub>, T<sub>a_out</sub>, VPD<sub>out</sub>, T<sub>a_in</sub>, T<sub>BS</sub>, T<sub>dry</sub> | Stepwise: index + env |
| 0.60 | 0.53 | 36 | CTD<sub>out</sub>, ICWSI<sub>in</sub>, CWSI<sub>dev_out</sub>, CSI<sub>out</sub>, CWSI<sub>Ta_in</sub>, TR<sub>out</sub>, VPD<sub>out</sub>, T<sub>BS</sub>, 502 nm, 520 nm, 576 nm, 727 nm, 746 nm, 766 nm, 823 nm, 920 nm, 979 nm, 1295 nm, 1407 nm, 1445 nm, 1519 nm, 1557 nm, 1893 nm, 1930 nm, 2451 nm, 2525 nm, 2042 nm | Stepwise: index + env + hyp |
| Lasso |  |  |  |  |
| 0.47 | 0.63 | 32 | CTD<sub>out</sub>, CSI<sub>in</sub>, CSI<sub>out</sub>, ICWSI<sub>in</sub>, ICWSI<sub>out</sub>, CWSI<sub>dev_in</sub>, CWSI<sub>dev_out</sub>, CWSI<sub>dev_gz</sub>, CWSI<sub>Ta_gz</sub>, T<sub>BS</sub>-T<sub>p</sub>, TR<sub>in</sub>, TR<sub>gz</sub>, T<sub>a_out</sub>, PAR<sub>gz</sub>, T<sub>BS</sub> | Lasso: index + env |
| 0.49 | 0.69 | 33 | CTD<sub>in</sub>, CTD<sub>out</sub>, CSI<sub>in</sub>, ICWSI<sub>in</sub>, ICWSI<sub>out</sub>, CWSI<sub>Ta_gz</sub>, T<sub>dry</sub> - T<sub>p</sub>, T<sub>a_out</sub>, VPD<sub>in</sub>, PAR<sub>gz</sub>, 403 nm, 449 nm, 468 nm, 904 nm, 1207 nm, 1213 nm, 2346 nm, 2377 nm, 2395 nm, 2426 nm, 2439 nm, 2445 nm, 2457 nm, 2476 nm, 2482 nm, 2525 nm | Lasso: index + env + hyp |
| 0.64 | 0.48 | 41 | T<sub>p</sub>, 492 nm, 495 nm, 526 nm, 545 nm, 576 nm, 604 nm, 610 nm, 626 nm, 645 nm, 654 nm, 658 nm, 667 nm, 724 nm, 1002 nm, 1006 nm, 1251 nm, 1276 nm, 1382 nm, 1395 nm, 1526 nm, 1893 nm, 2383 nm, 2426 nm, 2457 nm, 2476 nm, 2513 nm | Lasso: T<sub>p</sub> + hyp |
| 0.59 | 0.48 | 40 | 492 nm, 508 nm, 545 nm, 576 nm, 658 nm, 1238 nm, 1276 nm, 1395 nm, 1887 nm, 2506 nm, 2513 nm | Lasso: hyp |
| Random Forest |  |  |  |  |
| 0.54 | 0.57 | 33 | CTD<sub>out</sub>, CSI<sub>in</sub>, CSI<sub>out</sub>, ICWSI<sub>in</sub>, CWSI<sub>dev_in</sub>, CWSI<sub>dev_out</sub>, CWSI<sub>dev_gz</sub>, CWSI<sub>Ta_gz</sub>, T<sub>BS</sub>-T<sub>p</sub>, TR<sub>in</sub>, TR<sub>gz</sub>, T<sub>a_out</sub>, VPD<sub>gz</sub>, PAR<sub>gz</sub>, T<sub>BS</sub> | RF: index + env |
| 0.59 | 0.61 | 33 | CTD<sub>out</sub>, T<sub>dry</sub>-T<sub>p</sub>, T<sub>BS</sub>, T<sub>a_out</sub>, T<sub>dry</sub>, VPD<sub>out</sub>, PAR<sub>gz</sub>, 486 nm, 2488 nm, 2494 nm, 2500 nm, 2506 nm, 2513 nm, 2519 nm, 2531 nm | RF: index + env + hyp |
| Partial least square |  |  |  |  |
| 0.54 | 0.62 | 34 | CTD<sub>out</sub>, TR<sub>out</sub>, CWSI<sub>Ta_out</sub>, T<sub>BS</sub>-T<sub>p</sub>, CWSI<sub>dev_out</sub>, CSI<sub>out</sub>, T<sub>dry</sub>-T<sub>p</sub>, CWSI<sub>Ta_gz</sub>, CTD<sub>in</sub>, CWSI<sub>in</sub>, ICWSI<sub>out</sub>, TR<sub>in</sub>, ICWSI<sub>in</sub>, CWSI<sub>Ta_in</sub>, PAR<sub>gz</sub>, T<sub>BS</sub>, T<sub>a_out</sub>, T<sub>dry</sub>, T<sub>a_in</sub>, T<sub>a_gz</sub>, VPD<sub>out</sub>, VPD<sub>in</sub>, VPD<sub>gz</sub>, RH<sub>out</sub>, RH<sub>in</sub>, T<sub>p</sub>, 483 nm, 486 nm, 489 nm, 492 nm, 495 nm, 502 nm, 505 nm, 508 nm, 511 nm, 514 nm, 517 nm, 2488 nm, 2494 nm, 2506 nm, 2500 nm, 2513 nm, 2519 nm, 2525 nm, 2531 nm, | PLSR: index + env + hyp |

### Combining thermal infrared with hyperspectral data improves the prediction accuracy of the B104 transpiration model

While thermal infrared imaging informs on plant water-use behavior, hyperspectral reflectance spectra have been related to traits such as pigment content, photosynthetic efficiency, leaf anatomy and water content (67–69). Combining both may improve the accuracy and robustness of E prediction models, as significant correlations between hyperspectral data and g_s_ or E were observed (31, Fig. 5C). The wavelength regions that correlated with E were located around 523 nm, 992 nm, 1407 nm, 1881 nm and 2346 nm. All wavelengths, except for 523 nm, are related to water content and leaf anatomy (67,70). Six out of 10 selected indices (Table 1) correlated significantly with E (Fig. 5C, P<0.05). The strongest negative relationship was present between E and the WBI index (r=-0.52), while the most prominent positive correlation was observed with RMP_1483/1430_ (r=0.53). Both of these indices were created to monitor water content and a correlation between WBI and E has previously been demonstrated by Marino et al. (71). Adding hyperspectral indices (WBI and RMP_1483/1430_) to TIR index-based models significantly improved the CTD_out_ and CWSI_Ta_out_ models with ± 11% and ± 9%, respectively (Table 4, Fig. 6, Chi-square test, P<0.05). The absence of improvement in the T_dry_-T_p_ model may be related to the limited amount of hyperspectral information contained in these indices, a constraint that was reduced by training machine learning models (PLSR, RF, lasso, step-wise selection) on the complete hyperspectral, environmental and TIR datasets. The lasso and RF models with hyperspectral data were compared to the previously developed lasso and RF E prediction models, to evaluate the contribution of this imaging system. Adding hyperspectral data to E prediction models improved prediction accuracy to a small extent. The test RMSE of the RF model was slightly reduced by 10%, while the test R^2^ of both models was improved by 8 ± 2% (Table 5, Fig. 6). The best performing model was the lasso model, which had a RMSE of 0.49 and a R^2^ of 0.69. All algorithms selected environmental and thermal data, as well as hyperspectral information. The T_BS_, CTD_out_ and reflectance around 500 nm were selected by all model algorithms, while PAR, VPD_out_, T_dry_ – T_p_, ICWSI_in_ and reflectance around 2500 nm were predictors in three out of four models. All the variables selected by the models are listed in Table 5. The wavelengths selected by the E models were mainly related to water content and leaf anatomy (around 2500 nm) or photosynthesis/pigment content (around 500 nm). The hyperspectral models also included environmental data from all three monitoring positions directly or indirectly. Including environmental data into the E prediction models was pivotal as lasso models created using only image-based data (hyperspectral and/or T_p_) had approximately 30% lower prediction accuracy than the lasso model that incorporated all available data (Table 5).

### Validating drought detection and transpiration rate models on other maize genotypes

All empirical E prediction models developed so far were trained on the B104 genotype. The transferability of these models to other genotypes (H99, MS71, NC358, OH43, TX303, TZi8 and W153R) was uncertain, as genotypes may differ in drought sensitivity, water-use behavior, leaf anatomy and reflectance. In this study, all genotypes showed a similar drought response in measured physiological traits with larger treatment differences around the acute drought period, whereas the magnitude of these drought stress effects differed among genotypes (Fig. 8). Leaf ψ and ϕ_ps2_ showed the largest genotypic differences with more pronounced treatment effects in NC358, OH43, TZi8 and W153R compared to B104, H99, MS71 and TX303. However, only TX303 differed significantly from NC358, OH43 and W153R at 10 days after the onset of drought.

**Figure 7:**
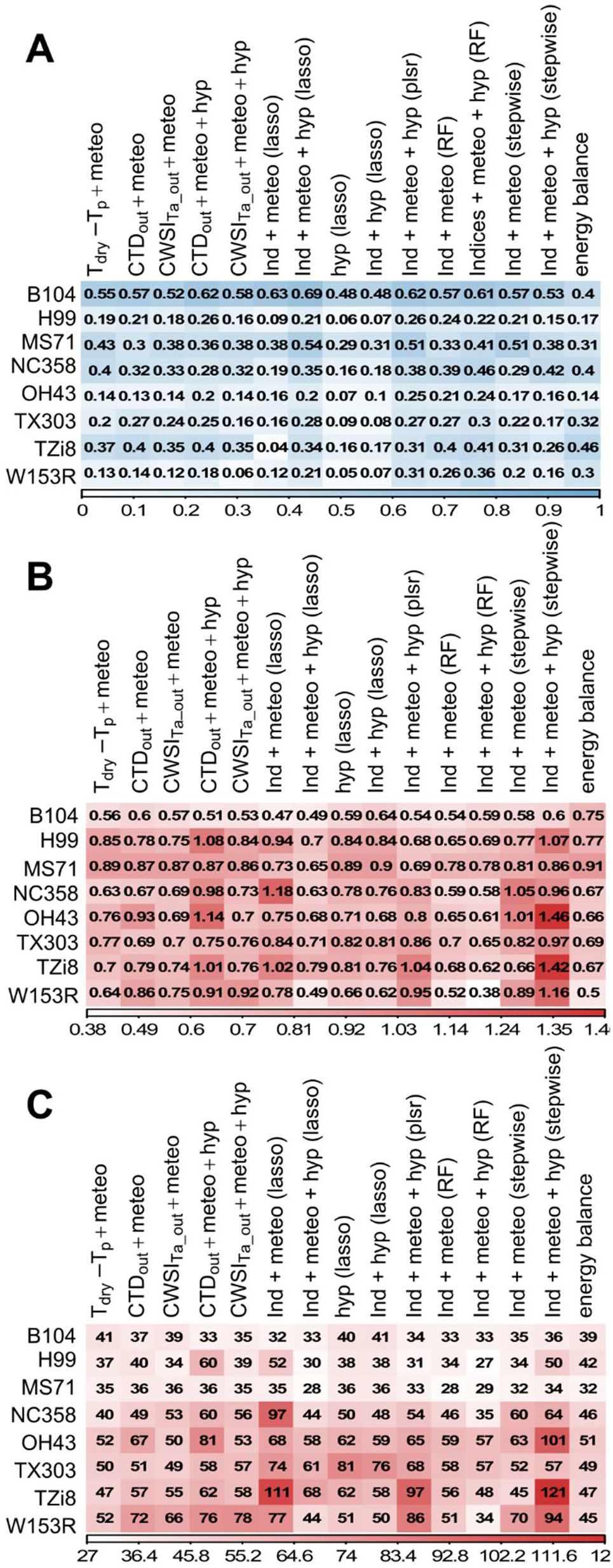
Accuracy of transpiration rate prediction models for different genotypes. Eight genotypes were compared: B104, H99, MS71, NC358, OH43, TX303, TZi8 and W153R. The test prediction accuracy of the 100 bootstrap samples was used for B104, while for the other genotypes the accuracy was determined with a test set that was not included in the training of the models. The prediction accuracy shown in this figure are (**A**) the R-squared (R^2^) colored in blue, and (**B**) the root-mean square error (RMSE) and (**C**) the mean absolute percent error (MAPE) colored in red. The shading in this figure represents the value of the accuracy measure with darker colors corresponding to higher values. Dark blue colors represent a high accuracy, while dark red colors indicate a low accuracy.

**Figure 8:**
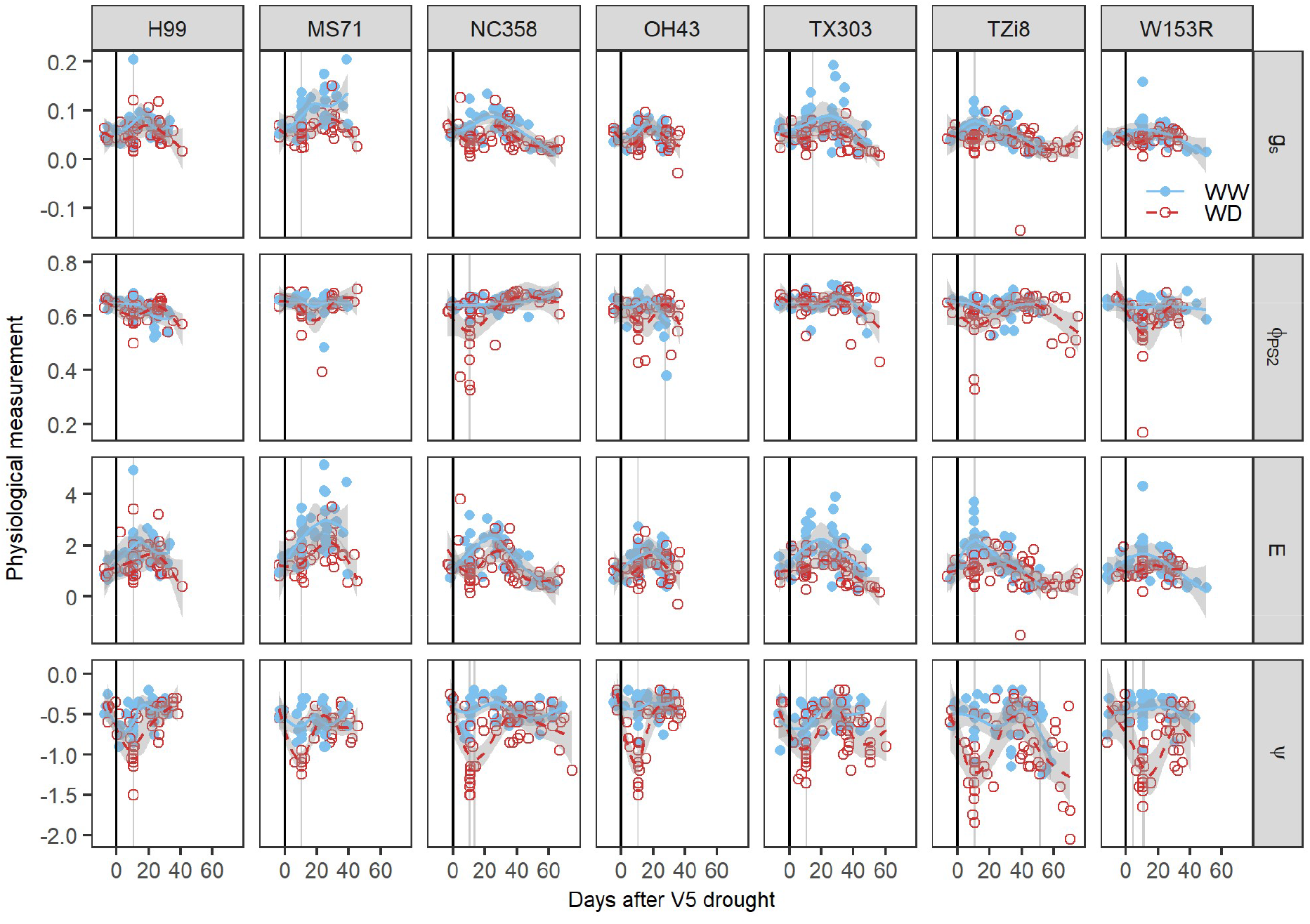
Genotypic differences in the responses of physiological traits to drought stress. The genotypes H99, MS71, NC358, OH43, TX303, TZi8 and W153R and four physiological traits, stomatal conductance (g_s_, mol H_2_O m^−2^s^−1^), effective quantum yield of photosystem II (φ_PS2_), transpiration rate (E, mmol H_2_O m^−2^s^−1^) and leaf water potential (ψ, MPa), were selected for this analysis. Physiological traits of well-watered (WW) and water-deficit (WD) maize plants were monitored from V4 (four fully developed leaves) until the silking stage. The average trends of the WW and WD treatments are indicated by blue solid and red dashed lines, respectively. The grey shading around the lines represents the 95% confidence interval of the average. Individual measurements of the plants are visualized by blue dots (WW) and red circles (WD). The black vertical line indicates the start of the drought treatment. The days on which significant treatment differences were observed are marked with a light gray vertical shading behind the average trends and dots (P<0.05).

Drought stress effects on E and g_s_ were less variable between genotypes. Treatment differences were only slightly larger in B104, MS71, NC358, TX303 and TZi8 compared to H99, OH43 and W153R (Fig. 8). This difference was mainly caused by higher E and g_s_ values in WW plants, while the treatment differences in ψ and ϕ_ps2_ resulted from decreases in the WD plants. The most drought-sensitive indices in B104 (CSI_in_ and TRI_in_) were able to detect drought stress effects in all eight genotypes and showed differences between genotypes. TX303 and TZi8 had the strongest drought stress effects, as significant treatment differences were observed during 28.5 ± 1.5 and 25.5 ± 1.5 days of the TF experiment, respectively. They were followed by B104, MS71 and NC358 with 20.5 ± 1.5, 20.5 ± 0.5 and 20.5 ± 0.5 days, respectively. The weakest drought stress effects were observed in H99, OH43 and W153R where only 15.5 ± 0.5, 8.5 ± 0.5 and 10 ± 1 days of the experiment showed significant treatment differences. The weaker drought stress effects in CSI_in_ and TRI_in_ of H99, OH43 and W153R compared to the other genotypes seemed to correspond with the genotypic differences observed in E and g_s_ (Fig. 9).

**Figure 9:**
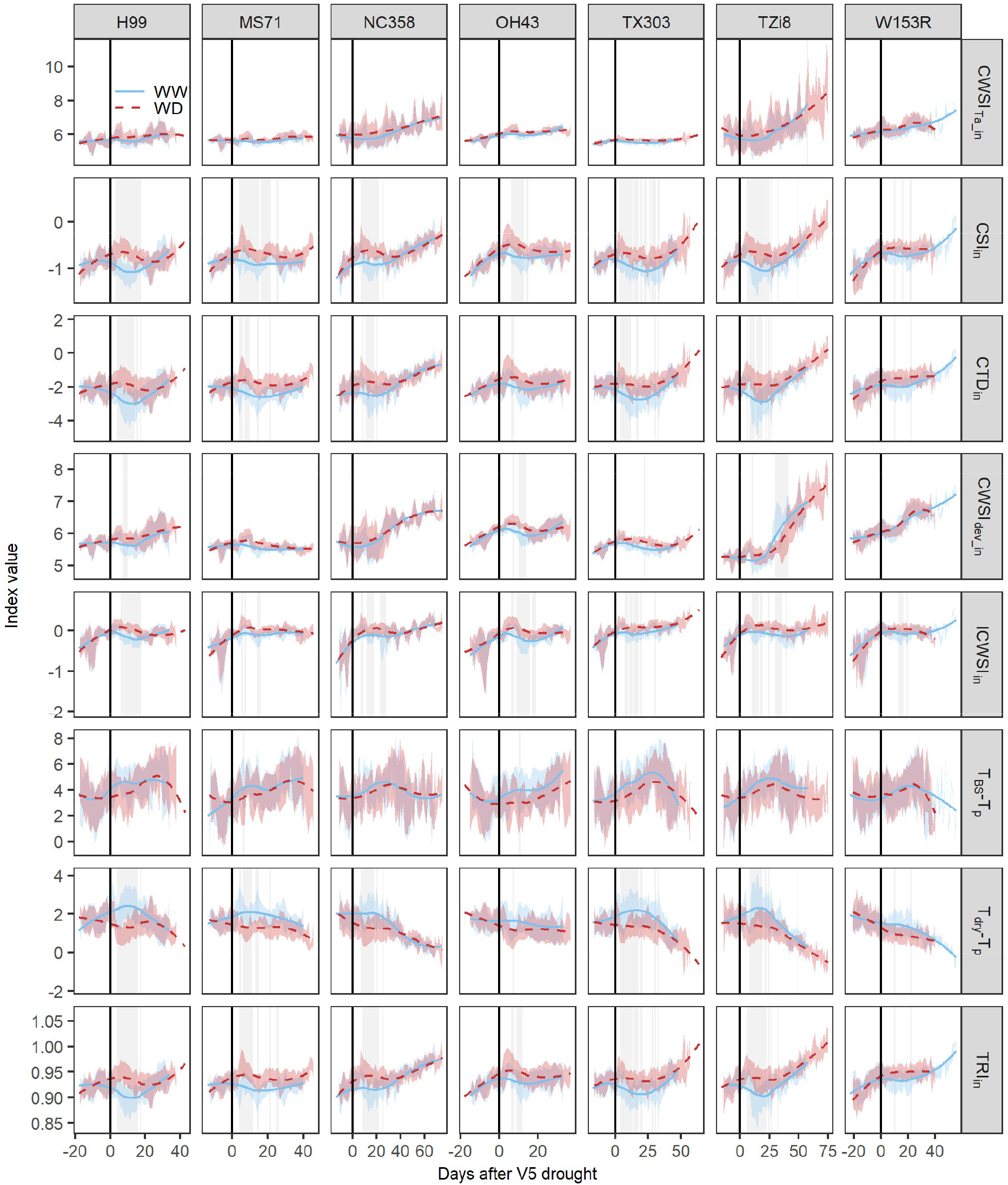
Responses of thermal infrared (TIR) indices to drought. The H99, MS71, NC358, OH43, TX303, TZi8 and W153R maize plants were imaged daily and TIR indices, including canopy temperature depression (CTD_in_), canopy stress index (CSI_in_), simplified stomatal conductance index (T_dry_-T_p_), temperature ratio index (TRI_in_), Idso’s crop water stress index (ICWSI_in_), crop water stress index (CWSI_Ta_in_, CWSI_dev_in_) and T_BS_-T_p_, were calculated using the formulas described in Table 1. The averages of the well-watered (WW, n: 19) and water-deficit (WD, n: 19) treatments are indicated by blue and red dashed lines, respectively. The blue and red shading represents the standard deviation around the mean. The black vertical line indicates the start of the drought treatment. The grey vertical shading indicates the days on which significant treatment differences were observed (P<0.05).

The relationship between E and TIR indices across genotypes influenced the prediction accuracy of the E prediction models. Most models were able to make reasonable predictions of E for multiple genotypes, except for the lasso model trained on TIR and environmental data, and the stepwise selection model that combined the environmental information with both hyperspectral and TIR data (Fig. 7). These models had small R^2^ (<0.1), high RMSE (>1) and MAPE (>90) values for certain genotypes. The high MAPE value indicated that the prediction error was higher than 90% of the measured value. The lasso and stepwise selection model had an acceptable prediction accuracy for B104, suggesting that they are not transferable to other genotypes. When comparing the remaining models, the RF, lasso and PLSR algorithms performed better than the other modelling approaches for most genotypes (Fig. 7). As the RF algorithm was trained with and without hyperspectral data, the contribution of this data type to the robustness and transferability of E prediction models could be evaluated. Adding hyperspectral data slightly improved the prediction accuracy for five out of eight genotypes, suggesting that the additional information contained in these data might improve the robustness of E predictions (Fig. 7). The RF model that combined both imaging data (TIR and hyperspectral) was further used to compare the prediction accuracy of different genotypes. The R^2^ of all seven genotypes was lower than the R^2^ of B104; however, the RMSE and MAPE values were more similar (Fig. 7). The lower R^2^ values may result from the larger E range in B104 compared to most other genotypes (Fig. 10). The genotypes could be subdivided into three groups based on the performance of the RF E model. The first group contained two genotypes, NC358 and W153R, for which the E model performed similar to B104 with an error of approximately 34% of the measured value (MAPE: ± 34% and RMSE>0.59, Fig. 7 and 10). The second group was more variable containing both low and high prediction errors. H99 had a low MAPE of only 22%, but a high RMSE of 0.69, which may be caused by the lower prediction accuracy of the higher E values (Fig. 10). In contrast, TZi8 and OH43 had a relatively high MAPE value (TZi8: 48%, OH43: 57%) combined with a relatively low RMSE value (TZi8: 0.62, OH43: 0.61). Figure 10 shows that for TZi8 and OH43 the E model slightly overestimated the measured E values. The last group contained MS71 and TX303 and showed the highest variability in the scatterplot comparing predicted to measured E values (Fig. 10). The prediction accuracy of these genotypes was low with an RMSE of 0.78 and a MAPE of 29% for MS71 and an RMSE of 0.65 and MAPE of 57% for TX303.

**Figure 10:**
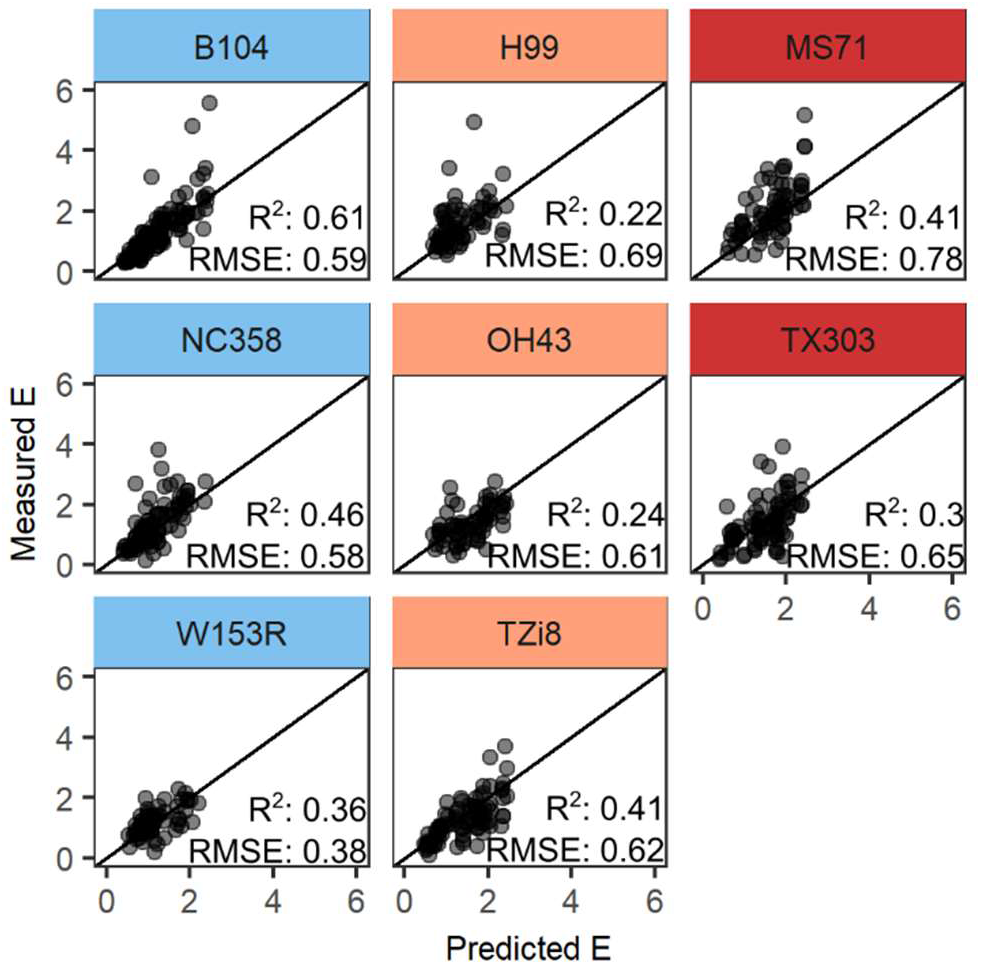
Relationship between measured and predicted transpiration rate (E) for different genotypes. The Random Forest models that combined thermal, hyperspectral and meteorological data were used to predict E. Genotypes were grouped based on the prediction accuracy of the model. The accurate group contained three genotypes (B104, NC358 and W153R) and is indicated by a light blue header. The moderately accurate group has an orange header and contains H99, OH43 and TZi8, while the less accurate group is marked with a red header and contains MS71 and TX303. The dots represent the predicted and measured E of individual plants, while the black line shows the one-to-one relationship. For B104, leave-one-out cross-validation predictions are shown, while the predictions of the other genotypes were created using a test set.

## Discussion

### Thermal infrared indices allow the detection of drought in an automated phenotyping platform

The use of TIR indices to detect drought is well established under field and orchard conditions, as this approach has been applied in irrigation/drought studies of potatoes, sesame, cotton, coffee, wheat, maize, forage grasses, pistachio, olive and peach (6,11,15,18,72–77). So far, the use of these indices is less common in indoor whole-plant phenotyping platforms. The few studies that used TIR indices in greenhouse experiments (6,18,76,78) showed that both the direct and empirical CWSI approaches, the I_g_ and CTD indices were able to detect drought in sesame, wheat, maize and peach trees. In this study, the CWSI index was not able to detect mild drought differences and its low drought sensitivity may be related to a lower accuracy of the non-water stressed baseline, as more variation was observed around the regression of VPD with CTD compared to the one with T_p_ (adj. R^2^_CTD_in-baseline_: 0.51, adj. R^2^_Tp_in-baseline_: 0.66, Additional file: Fig. S1). The baseline relating VPD to T_p_ was used to calculate the ICWSI index, which was, in contrast to the CWSI, able to detect treatment differences eight days after the onset of drought. Increasing the sample size used to develop the CWSI baseline may improve the performance of this index. In addition to the lower accuracy of the baseline, inaccurate estimations of the CTD of a non-transpiring plant (3 °C) may reduce the drought sensitivity of the CWSI. Alternative methods to estimate this CTD value include measuring T_a_ and T_p_ of a leaf covered with petroleum jelly or calculating CTD based on the difference between the saturated vapor pressure at T_a_ and at the temperature of a non-transpiring plant, which can be estimated by adding T_a_ to the intercept of the relationship between VPD and CTD of a fully transpiring plant (6,25,79). On the other hand, the drought sensitivity of the CTD_in_ index was confirmed in this study, as this index detected significant drought stress effects four days after the onset of drought. Nevertheless, it was outperformed by the CSI_in_ and TRI_in_ indices, which showed significant differences on the second day. The improved performance of the CSI_in_ index may result from the diurnal VPD trend that was present in the greenhouse. This VPD range created a larger range in E and consequently T_p_ values which could not be normalized by T_a_ alone, but required a correction for RH (44). The CSI_in_ index incorporates a correction for RH by normalizing CTD with VPD. The CSI was originally developed to detect drought stress in wheat fields. Rodriguez et al. (48) observed that this index captures physiological responses both at the single-plant leaf and population-field level.

### Transpiration rate can be predicted based on thermal infrared indices or the energy balance approach

Plant temperature and TIR indices have been related to different water-use behavior traits, such as g_s_, E, relative water content, and stem and leaf water potential (11,12,14,18,22,27,28,76,80). One of the drought-sensitive indices in this study, CSI, has been correlated with g_s_ and E in the field (48), but this was not confirmed in this study under greenhouse conditions. CSI_in_ was not correlated with the gas-exchange measurements, while CSI_out_ was non-linearly related to these traits. A negative relationship of CSI_out_ with g_s_ and E was observed when the index values exceeded −1.5, which corresponds with the results observed by Rodriguez et al. (48). Points that deviated from this negative relationship (CSI_out_ < −1.5) had E values <2 mmol m^−2^s^−1^ and were collected in the morning when the stomata were not completely open yet (Additional file: Fig. S3). The opening of the stomata is triggered by light (81) and it requires some time for all the stomata to open and for E to reach its maximal value. During this time period the relationship between VPD and T_p_ is less strong or even absent (44), which may explain the lower performance of the CSI_out_ index during these time points. The TIR indices CTD, CWSI_Ta_ and T_dry_-T_p_ (R^2^_CTD_out_: −0.63 – −0.64, R^2^_CWSI_Ta_out_: −0.58 – −0.61, R^2^_Tdry-Tp_: 0.5 – 0.61) did have a relatively strong correlation with the gas-exchange measurements. Negative relationships between CTD or CWSI and g_s_ have been observed in olive trees, sesame, strawberry, grapevine, both in field and greenhouse setups (12,18,22,27,82). Maes and Steppe (25) simulated the relationship between g_s_, CTD and the direct CWSI approach for different environmental conditions and observed a non-linear relationship between these indices and g_s_. However, several studies have described a linear or almost linear relationship of CTD and CWSI with g_s_ (18,22,51,82). Some of these more linear relationships may have resulted from a low g_s_ range, as g_s_ is almost linearly related with T_p_ and CWSI as long as its value is smaller than <0.5 mol m^−2^s^−1^ (17). The other strongly correlated index T_dry_-T_p_ had a positive relationship with g_s_ and E. It is a simplified version of the stomatal conductance index (I_g_,(51)) and describes an almost linear relationship between this index and g_s_, which was also visible in this study (Additional file: Fig. S3).

The relationship between E or g_s_ and TIR indices can be used to develop empirical prediction models, but only a few studies have used this to predict physiological traits such as g_s_, and stem and leaf ψ (11,26,83,84). The limited use of empirical models may result from the effects that environmental factors, such as T_a_, PAR, wind speed and VPD, have on the relationships between these water use behavior traits and TIR indices (25), which, in the field, may differ within and between days reducing the reliability of these models. On the other hand, the environmental conditions in indoor phenotyping platforms are semi-controlled, making the use of these types of models more feasible. In this study, TIR index-based models had higher prediction accuracies when indices were combined with environmental data, such as VPD and PAR. The best performing model was the RF model that incorporated TIR indices and environmental variables collected at the three monitoring positions directly or indirectly. The need to monitor the environment at different positions in the greenhouse resulted from the fact that the plants were moved during the imaging cycle. The prediction accuracy could probably be further improved by incorporating the exact position of the plant in the growth zone and interpolating the environmental conditions for these positions. Environmental monitoring outside the growth zone of the platform is not required in phenotyping platforms with mobile imaging systems in which the cameras move to the plants. However, in these platforms imaging is performed in a less controlled light environment and plants are often grown in a canopy structure to simulate field conditions, which will increase the influence of shading on T_p_ and E predictions. Within-canopy illumination variation will induce leaf temperature variation that is unrelated to changes in g_s_ and E leading to scatter in the canopy temperature-physiology relationships. To reduce the non-biological variation in temperature, the more sensitive sunlit pixels can be extracted from the images (85,86), which have a more pronounced slope between CTD and g_s_ (25,87), but can have a higher variability in T_p_ and g_s_ that may mask subtle differences in the plant water status (86,88). More research is needed to elucidate the effects of illumination variation in indoor thermography applications and develop optimal approaches that can remove these effects.

Empirical models are platform-specific and cannot be transferred to the field, whereas the energy balance approach does not have this limitation. This mechanistic model has been evaluated in field studies (29,89,90), but its use in indoor phenotyping platforms is less common because of the difficulty of distinguishing the longwave radiation from the surrounding environment (91). The need of measuring net isothermal radiation can be eliminated by using a simplified version of the energy balance approach that incorporates the temperature of a non-transpiring leaf (dry reference surface). This simplified energy balance model was evaluated in this study, however, the approach had a lower accuracy compared to the empirical models due to the incorrect model assumptions (Figure 7). A detailed discussion of the issues observed during the implementation of this mechanistic model and suggestions on how to optimize this can be found in the additional file: file S1. One important aspect of the energy balance approach is the steady-state assumption, which states that the environmental conditions are stable during imaging and that leaf temperature is at equilibrium with its environment. This assumption is difficult to maintain in the field as the environmental conditions are changing during the day. In indoor phenotyping systems environmental conditions are often semi-controlled making it easier to monitor the plants in steady-state conditions. However, there are still factors, such as plant transportation, changing light levels, and diurnal temperature and VPD trends, that may influence the acclimatization of the plants to the imaging environment and the steady-state condition. Vialet-Chabrand and Lawson (91) proposed a solution to this problem by developing a method that can predict g_s_ and E under dynamic environmental conditions. This is accomplished by applying an energy balance model with an in-built dynamic model of g_s_, which is fitted on observed temperature measurements. This new method is able to derive g_s_ under a fluctuating light environment and detects temporal response variations within and between leaves (17,91).

### Hyperspectral data may improve the robustness of E prediction models

Hyperspectral reflectance in the near-infrared (NIR, 700-100 nm) and short-wave infrared (SWIR, 1000-2500 nm) region has been related to leaf cuticle thickness, water content and anatomy (70,92). These traits can influence E directly or indirectly by determining total leaf conductance, which is influenced by g_s_, cuticular transfer, intercellular space and mesophyll cell wall conductance (1). Many methods that estimate g_s_ provide a measurement of total leaf conductance, which is most strongly affected by g_s_. The relationship between g_s_ or E and hyperspectral reflectance was confirmed in this study and in Mertens et al. (31). Both studies showed significant relationships with blue-green (523-532 nm) and the NIR water-absorption trough (976-992 nm) reflectance. The number of significant correlations was higher in this study, as additional wavelengths located in the SWIR water absorption troughs (1407, 1881 and 2346 nm) also correlated with E and g_s_. The diurnal relationship between red reflectance and E was the only one described by Mertens et al (31) that wast not detected here. The reason may be the limited number of E measurements collected within one day and/or the interaction between drought and diurnal effects, which increased the changes in red reflectance and decreased E variation. Overall, several studies have used the relationships between reflectance and E to develop hyperspectral indices that can be used to monitor this trait (37,52,71). The WBI index was originally created to monitor water content in plants, but Marino et al. (71) observed correlations with g_s_ and whole-plant E, which were also visible in this study (r_WBI-E_ = −0.52). This correlation differs from the results observed by Mertens et al. (37); where WBI was not significantly related to E. These differing results may be caused by dissimilarities in the time between imaging and measurements, which ranged up to 1 hour in the study of Mertens et al. (37) and was limited to 10 min in this study. The other two indices that had relatively high correlations with E were SR_1440/1460_ (r=-0.49) and RMP_1483/1430_ (r= 0.53). These indices included (SWIR) water absorption regions, suggesting that these parts of the spectrum may contain information about E.

The combination of thermal and hyperspectral indices in empirical models has mainly been used to predict yield, chlorophyll content and relative water content (14,32). These studies observed a slight improvement in R^2^ when thermal and hyperspectral indices were combined in a PLSR or stepwise multiple linear regression model. Adding hyperspectral information to predict E was only beneficial here for the single TIR index-based models that used CWSI_Ta_ and CTD. In these models, the water content WBI and RMP_1483/1430_ indices replaced the VPD variable suggesting that they may contain additional information about the relationship between leaf water content, E and VPD. When hyperspectral reflectance was combined with multiple TIR indices and environmental data a slight improvement was visible; however, this combination performed better for some of the other inbred lines, especially W153R. This improvement suggests that hyperspectral data can capture genotypic differences in water use behavior, but more research is needed to determine the contribution of this data type to E prediction models. The most robust approach to predict E was the energy balance model (Additional file: file S1), which had similar, albeit low (R_2_=0.32, RMSE=0.68 and MAPE=45%) prediction accuracies for all eight inbred lines. The accuracy of this approach can be improved by refining the T_dry_ estimate, which may make this a viable method to monitor E in automated phenotyping platforms.

### TIR indices can detect genotypic differences in drought sensitivity

Genotypic differences in plant temperature or TIR indices have been observed in several heat tolerance or drought sensitivity studies on rice, wheat and maize (16,23,93–95). These studies noted larger CWSI differences between WW and WD treatments in drought stress-sensitive genotypes compared to tolerant ones. In this study, genotypic differences in drought sensitivity were observed in both TIR indices and physiological traits. The strongest physiological differences were present in the leaf ψ measurements in which the drought induced decrease was significantly less pronounced in TX303 than OH43, NC358 and W153R. This partially corresponded with the available drought sensitivity data in which plant height and leaf length of OH43 and W153R were more affected by drought compared to TX303 (data not shown). These genotypic differences diverged from the CSI_in_ and TRI_in_ results in which TX303 was more affected by drought compared to OH43, W153R and H99. The index results seemed to correspond with the g_s_ and E measurements, which showed slightly larger (but not significant) treatment differences in TX303 compared to the other three genotypes. The limited number of significant differences observed in the genotype comparison analysis may have resulted from confounding factors. Measurements and images were grouped based on how long the plants had received the drought treatment, which started at the V5 stage. This procedure was chosen to facilitate the comparison of genotypes that differed in developmental timing and thus the date at which the drought started. This resulted, however, in combining and comparing data collected on different measurement days. The potential difference in environmental conditions may have created additional variation in the E and g_s_ measurements. To make this analysis more robust, a larger number of physiological E and g_s_ measurements are required, or the start of the drought treatment should be synchronized for all the genotypes. Consequently, more research is needed to elucidate the genotypic differences in physiology and TIR indices of these eight inbred lines.

## Conclusion

Thermal infrared imaging has been a popular tool to detect drought and estimate g_s_ or E in the field for many years. The use of thermography in indoor automated phenotyping platforms has been less investigated and is mainly limited to comparing T_p_ or CWSI between drought-stressed and well-watered plants. This study demonstrates the use of TIR indices, empirical and mechanistic models to detect drought and predict E of maize in an automated phenotyping platform. TIR indices that corrected T_p_ for VPD and/or T_a_ (CSI, TRI), were most sensitive to drought and were able to detect genotypic differences but were not strongly correlated with E and g_s_. Instead, E and g_s_ were correlated with the commonly used indices CWSI and CTD, which could be used to develop empirical E prediction models. Model performance was highest when thermal indices were combined with environmental data and hyperspectral indices or wavelengths. Empirical models had the highest prediction accuracies for the genotype it was trained on (B104), but their performances were inconsistent for other genotypes indicating that the genotypic differences should be considered during model development. In addition, it is important to monitor environmental data at multiple positions of a phenotyping platform with fixed camera positions, as transportation will influence the acclimatization of the plant to its environment and E prediction accuracy.

## Abbreviations

CSI: Canopy Stress Index
CTD: Canopy Temperature Depression
CWSI: Crop Water Stress Index
CWSI_dev_: CWSI corrected for development
CWSI_Ta_: CWSI corrected for non-constant air temperature conditions
DR: small drought experiment
dSR_660/1040_: 1st derivative Simple Ratio index of 660 and 1040 nm
E: transpiration rate
F_v_’/F_m_’: energy harvesting efficiency by oxidized PSII
g_s_: stomatal conductance
gz: growth zone
ICWSI: Idso’s Crop Water Stress Index
In: inside cabin
LED: light emitting diode
MAPE: mean absolute percent error
ND_1425/2145_: Normalized Difference index of 1425 and 2145
nm NDI_1407/1862_: Normalized Difference Index of 1407 and 1862 nm
NDWI: Normalized Difference Water Index
out: outside cabin
PAR: photosynthetically active radiation
PLSR: Partial Least-Squares Regression
R^2^: R-squared
R_953/492_: Ratio index of 953 and 492 nm
RF: Random Forest
RGB: red-green-blue
RH: relative humidity
RMP_1483/1430_: Relative Moisture Percentage of 1483 and 1430 nm
RMSE: Root Mean Square Error
SDD: Stress Degree Days
SR_1440/1460_: Simple Ratio index of 1140 and 1460 nm
T_a_: air temperature
T_BS_: black sphere temperature
T_dry_: dry reference leaf temperature
T_dry_-T_p_: simplified stomatal conductance index
TF: transferability experiment
TIR: thermal infrared
T_p_: plant temperature
TRI: Temperature Ratio Index
T_wet_: wet reference leaf temperature
VPD: vapor pressure deficit
WBI: Water Band Index
WCI: Water Content Index
WD: water-deficit
WPI_2_: Water Potential Index 2
WW: well-watered
ϕ_ps2_: effective quantum yield of photosystem II
ψ: water potential

## Acknowledgements

We wish to thank Drs. Liqun Xing and Erin Slabaugh for technical input.

## Declarations

### Ethics approval and consent to participate

Not applicable.

### Consent for publication

Not applicable.

### Availability of data and materials

The datasets generated and analyzed during the current study are available in the zenodo repository (https://doi.org/10.5281/zenodo.7807989, https://doi.org/10.5281/zenodo.8164473, https://doi.org/10.5281/zenodo.8033640)

### Authors’ Contributions

N.W., D.I., H.N., J.V., W.B, S.Ma conceived the original screening and research plans; L.V., H.S., S.Me., J.M. and B.C. performed the experiments, collected the thermal and hyperspectral data and physiological measurements; K.D. and J.D.B. provided technical assistance during experiments; S.D.M developed the RGB segmentation model; S.Me. analyzed the thermal and hyperspectral data, performed statistics and wrote the article; N.W., D.I. and S.Ma. revised the article. D.I. agrees to serve as the author responsible for contact and ensures communication.

## Authors’ information

### Current addresses

H.S. present address: German Federal Institute for Risk Assessment, Department Food Safety, Max-Dohrn-Str. 8-10, 10589 Berlin, Germany

J.M. present address: Instituut voor Landbouw, Visserij-en Voedingsonderzoek (ILVO), Eenheid Plant, Caritasstraat 39, 9090 Melle, Belgium

N.W. present address IBG-2: Plant Sciences, Forschungszentrum Jülich GmbH, 52425 Jülich, Germany

S.D.M present address: Robovision, Technologiepark 80, Zwijnaarde, Belgium

J.V. and W.B. present address: BASF Corporation, 2TW Alexander Drive, Research Triangle Park, North Carolina, 27709, U.S.A.

## Supplementary Information

### Additional file 1: Figure S1. PDF

Baselines of the crop water stress indices. This figure illustrates the baselines used to calculate the Idso crop water stress index (ICWSI), development-corrected crop water stress index (CWSI_dev_) and air temperature (T_a_) corrected crop water stress index (CWSI_Ta_) inside the imaging cabin. Similar baselines were created for the other monitoring positions (outside the cabin, growth zone). In A and B, the baselines of juvenile and mature plants are represented by a black and yellow line, respectively, while the individual measurements are indicated with black and yellow dots. **A**, baselines of the ICWSI_in_ index, which relates plant temperature (T_p_) to vapor pressure deficit (VPD). Separate baselines were created for the different genotypes and juvenile/mature plants. This baseline was used to estimate plant temperature (T_p_) of a fully transpiring plant. **B**, the baselines of the CWSI_dev_in_, which relates canopy temperature depression (CTD = T_p_-T_a_) to VPD. This function is used to estimate the CTD of a fully transpiring plant. Separate baselines for genotypes and developmental stages were also created for this index. **C**, representation of the baselines used to calculate the CWSI_Ta_in_. The model of this baseline has CTD as the dependent variable and VPD, T_a_ and its interaction term as independent continuous variables. This figure illustrates what the relationship between CTD and VPD would look like if T_a_ was constant. The relationships between CTD and VPD for eight different temperatures are indicated by solid lines. The measurements are visualized by slightly transparent dots. Each temperature has received a unique color, which is used for both the line and dots. Separate baselines were developed for each genotype.

### Additional file 2: File S1. PDF

Energy balance transpiration rate model. Detailed description of the simplified energy balance transpiration rate model formulas, results and critical discussion of the implementation on PHENOVISION.

### Additional file 3: Figure S2. PDF

Environmental data. The daily mean air temperature (Ta, **A**), relative humidity (RH, **B**) and vapor pressure deficit (VPD, **C**) of the three monitoring positions (gz, in and out) are represented by a solid light gray line, dashed dark gray line and dotdashed black line, respectively. **D** shows the measurements of the black sphere temperature (T_BS_), PAR monitored in the growth zone, and the dry reference temperature (T_dry_) that was measured inside the imaging cabin. The daily mean of T_BS_ and PAR are indicated by a thin and thick light gray line, respectively, while T_dry_ is represented by a dashed dark gray line.

### Additional file 4: Figure S3. PDF

Relationship between thermal infrared indices and transpiration rate (E). Individual measurements are represented with colored dots showing the VPD_out_ at the time of sampling. A blue-red gradient is used to visualize the time range. The linear or polynomial relationships between the indices are indicated with a black line, while non-linear (spline) relationships are represented by a blue line. The gray shading around the lines show the 95% confidence interval of the relationship. E (mmol m^−2^s^−1^) versus (**A**) CSI_out_, (**B**) CTD_out_, (**C**) CWSI_Ta_out_, and (**D**) T_dry_-T_p_ (°C).

