## Additional file 1: Figure S1. for "Monitoring of drought stress and transpiration rate using proximal thermal and hyperspectral imaging in an indoor automated plant phenotyping platform"

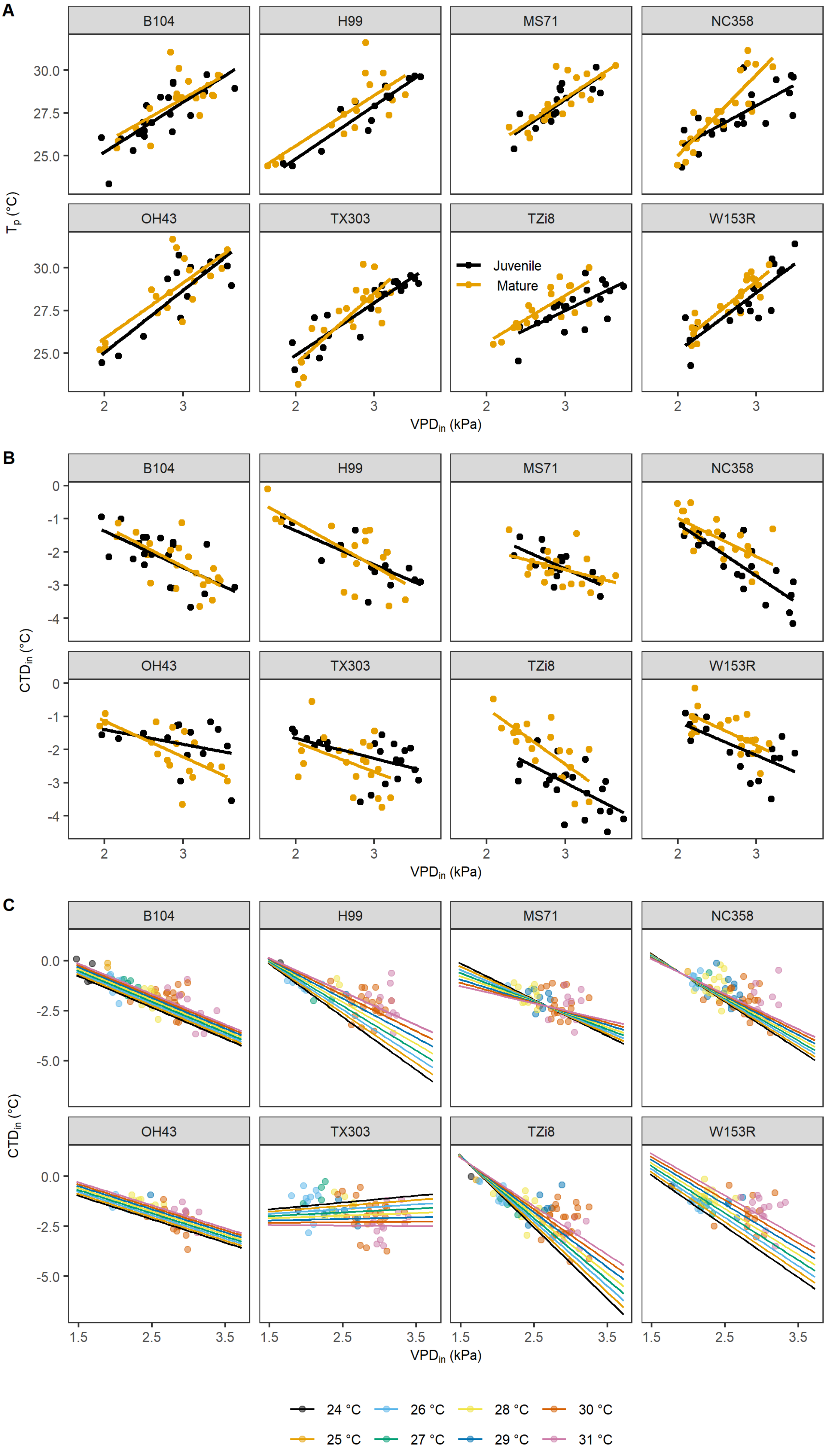


Baselines of the crop water stress indices. This figure illustrates the baselines used to calculate the Idso crop water stress index (ICWSI), development-corrected crop water stress index ($CWSI_{dev}$) and air temperature ($T_a$) corrected crop water stress index ($CWSI_{Ta}$) inside the imaging cabin. Similar baselines were created for the other monitoring positions (outside the cabin, growth zone). In A and B, the baselines of juvenile and mature plants are represented by a black and yellow line, respectively, while the measurements are indicated with black and yellow dots. **A**, baselines of the $ICWSI_{in}$ index, which relates plant temperature ($T_p$) to vapor pressure deficit (VPD). Separate baselines were created for the different genotypes and juvenile/mature plants. This baseline was used to estimate plant temperature ($T_p$) of a fully transpiring plant. **B**, the baselines of the $CWSI_{dev\_in}$, which relates canopy temperature depression (CTD = $T_p$-$T_a$) to VPD. This function is used to estimate the CTD of a fully transpiring plant. Separate baselines for genotypes and developmental stages were also created for this index. **C**, representation of the baselines used to calculate the $CWSI_{Ta\_in}$. The model of this baseline has CTD as the dependent variable and VPD, $T_a$ and its interaction term as independent continuous variables. This figure illustrates what the relationship between CTD and VPD would look like if $T_a$ was constant. The relationships between CTD and VPD for eight different temperatures are indicated by solid lines. The measurements are visualized by slightly transparent dots. Each temperature has received a unique color, which is used for both the line and dots. Separate baselines were developed for each genotype.
