## Additional file 2: File S1. for "Monitoring of drought stress and transpiration rate using proximal thermal and hyperspectral imaging in an indoor automated plant phenotyping platform"

### Simplified energy balance description, results and discussion

#### Methods

##### Energy balance transpiration rate model

The energy balance describes the theoretical relationship between surface temperature and evapotranspiration. This relationship is given by:

$$R_n = H + \lambda E + G_i + S \qquad (1)$$

in which $R_n$ represents the net radiation (W $m^{-2}$), H is the sensible heat flux (W $m^{-2}$), λE represents the latent heat flux or the energy transfer related to evapotranspiration (W $m^{-2}$), $G_i$ is the soil heat flux (W $m^{-2}$) and S represents the total aboveground energy storage (W $m^{-2}$). $G_i$ and S are generally small compared to the other terms and can often be ignored in the leaf energy balance calculations (1). The energy balance equation can be converted to a formula that describes the relationship between $T_p$-$T_a$ and the energy flux (2):

$$T_p - T_a = \frac{[r_{HR}\,(r_{aw}+r_s)\,\gamma\,R_{ni} - \rho\,c_p\,r_{HR}\,VPD]}{[\rho\,c_p(\gamma(r_{aw}+r_s)+s\,r_{HR})]} \qquad (2)$$

In this formula $T_p$ and $T_a$ (°C) are plant and air temperatures, $r_{HR}$ is the parallel resistance to heat and radiative transfer (s $m^{-1}$), $r_{aw}$ represents the boundary layer resistance to water vapor (s $m^{-1}$), $r_s$ is the leaf resistance to water vapor, which is assumed to be dominated by stomatal resistance (s $m^{-1}$), γ represents the psychometric constant (66.8 Pa $C^{-1}$, (3)), $R_{ni}$ is the net isothermal radiation (W $m^{-2}$), ρ is the density of air (1,204 kg $m^{-3}$, (3)), $c_p$ represents the specific heat capacity of air (1013 J $kg^{-1}$ $C^{-1}$, (3)), VPD is the air vapor pressure deficit (Pa) and s is the slope of the curve relating saturating water vapor pressure to temperature (Pa °$C^{-1}$, (4)). This equation can be converted to calculate the stomatal resistance $r_s$ or conductance $g_s$ ($g_s$=1/$r_s$) from $T_p$ and meteorological variables (2) :

$$r_s = -\rho\,c_p\,\frac{r_{HR}[s\,(T_p-T_a)+VPD]}{[\gamma(\rho\,c_p(T_p-T_a)-r_{HR}\,R_{ni})]} - r_{aw} \qquad (3)$$

This equation requires an estimate of $R_{ni}$ which was not available in PHENOVISION; however, the need of this variable can be eliminated by using the temperature of a dry reference leaf ($T_{dry}$, °C).

$$T_{dry} - T_a = \frac{R_{ni}\, r_{HR}}{\rho\, c_p} \quad (4)$$

$T_{dry}$ was acquired in this study by placing a black metal plate within the field of view of the camera. This black plate and subsequently $T_{dry}$ was measured together with each plant. By assuming the dry reference had similar aerodynamic and optical properties as the plant of interest equation 4 could be rearranged to (2):

$$r_s = -\rho\, c_p r_{HR} \frac{[s\,(T_p - T_a) + VPD]}{[\gamma\rho\, c_p(T_p - T_{dry})]} - r_{aw} \quad (5)$$

The variable $r_{HR}$ was estimated according to Maes and Steppe (1), while $r_{aw}$ was calculated as 0.92 * resistance to sensible heat ($r_{ah}$) (3).

$$r_{ah} = 0.5\left(r_{ah,u(forced)} + r_{ah,u(free)}\right)^{-1} \quad (6)$$

$$r_{ah,u(free)} = \frac{400}{abs\left(T_p - T_a\right)^{0.33}} \quad (7)$$

$$r_{ah,u(forced)} = 100\left(\frac{D}{u}\right)^{0.5} \quad (8)$$

$$D = \sqrt{\frac{A}{\pi}\left(\frac{1}{\frac{W}{L}} + \frac{W}{L}\right)} \quad (9)$$

In equation 6 $r_{ah,u(forced)}$ is the resistance of the air above the leaf surface to heat transfer through forced convection, while $r_{ah,u(free)}$ is the resistance of the air to sensible heat exchange through free air convection (1). $r_{ah,u(free)}$ can be calculated using equation 7 and $r_{ah,u(forced)}$ using equation 8. To estimate $r_{ah,u(forced)}$ the leaf/plant dimensions and wind speed have to be known. In this study the wind speed was assumed to be 0.01 m $s^{-1}$, which is a low value as there was almost no wind flow present in the greenhouse, while the area (A), width (W) and length (L) of the plant were extracted from top view RGB images.

To convert $g_s$ to E, the latent heat flux (λE) was calculated using equation 10 (1).

$$\lambda E = \rho\frac{c_p}{\gamma} * \frac{e_s^*(T_p) - e_a}{r_v} \quad (10)$$

In this equation $e_a$ is the vapor pressure in the air (kPa), $e_s^*(T_p)$ is the saturated vapour pressure at the plant surface and $r_v$ is the total resistance to vapor transport (s $m^{-1}$). To calculate $r_v$ the equation of isolateral leaves was used (equation 11) which assumed that the stomatal resistance ($r_s$) and the boundary layer resistance ($r_{aw}$) was the same for the upper and lower side of the leaves (1).

$$r_V = \frac{r_s + r_{aw}}{2} \tag{11}$$

This simplification was necessary as no information on the difference in stomatal density and boundary layer between the adaxial and abaxial side was available. To convert λE in (W $m^{-2}$) to transpiration rate (mmol $m^{-2}$ $s^{-1}$) the following equation was used:

$$E = \left[\frac{\frac{\lambda E}{l_w}}{M_w}\right] * 10^3 \tag{12}$$

with $l_w$ the latent heat of vaporization (J $g^{-1}$) and $M_w$ the molar mass of water in (g $mol^{-1}$).

The accuracy of the energy balance model was determined by comparing the E estimates with the infrared gas analyzer measurements of E. The MAPE, RMSE and $R^2$ values of the energy balance model were calculated with equation 2 and the 'postResample' function ( 'caret' R package, (5)).

### Results

#### Transpiration rate can be predicted using empirical and mechanistic modelling approaches

The models discussed in the main article are empirical models that are suited to estimate E in PHENOVISION, but cannot be transferred to other setups, as the relationships and importance of predictors may differ. This issue can be solved by calculating transpiration rate using the energy balance method. Stomatal conductance and subsequently transpiration rate can be directly derived from plant temperature and environmental variables, such as net isothermal radiation, wind speed, Ta and RH (1,2). Net isothermal radiation is not measured in PHENOVISION; however, the need of this variable can be avoided by using the temperature of a dry reference leaf. In this study, a black metal reference plate was imaged together with the plant ($T_{dry}$) and used to test the energy balance approach by assuming it had similar aerodynamic and optical properties as the plant of interest. The environmental data outside the cabin was selected for this approach, as this was the environment at which E was measured. The performance of the energy balance model was lower than the empirical models (RMSE=0.75 and $R^2$=0.40, Table 6, Fig. 7), However, it was less sensitive to genotypic differences, compared to the best performing random forest model that combined thermal infrared, hyperspectral and environmental data. The median $R^2$ value for all genotypes was 0.32, while the median RMSE and MAPE were 0.68 and 45%, respectively. Overall, there appears to be a trade-off between the accuracy and the robustness of the energy balance model.

### Discussion

#### Transpiration rate can be predicted based on the energy balance approach

In this study, the accuracy of a simplified energy balance model was found to be much lower compared to the accuracy of the empirical RF model. The reason lies with the estimate of $T_{dry}$, which should be determined using a reference material that has the same optical and thermal properties as the leaves (6). The reference used in this study strongly differed from maize leaves, as it was a relative thick black metal plate positioned horizontally in the imaging cabin with the purpose to check for camera temperature drift. The $T_{dry}$ estimation could be improved by determining the relationship between the temperature of the reference plate and a non-transpiring leaf covered in petroleum jelly. Such correction may still introduce some error, as adding jelly to a leaf might alter its properties including its emissivity. $T_{dry}$ estimates can also be improved by calculating the temperature of a non-transpiring leaf using an energy balance model that incorporates the difference in optical and thermal properties between the reference and the leaf (7). This approach will give the most accurate estimation of $T_{dry}$, but it requires additional information on the properties of the material. Another factor that may have influenced the accuracy of the energy balance model, is the fact that the plants were not acclimatized to the environmental conditions in which $T_{dry}$ was measured. In Phenovision the dry reference is installed inside the imaging cabin, while the plants are immediately imaged upon entering the cabin causing them to be adjusted to the growth zone environment instead. This setup was implemented to avoid the acclimatization of plants to irrelevant environmental conditions during imaging, however, it can complicate the implementation of the energy balance approach. This issue can be reduced by implementing the previously described $T_{dry}$ estimation improvement, namely determining the relationship between reference plate temperature and non-transpiring leaf temperature of a plant that was transported to the imaging cabin. Within-plant variations in $g_s$ and E could also have affected the accuracy of our protocol, as point measurements were used to validate plant average E predictions. It would have been better to predict E for the measurement position or to compare it to an average of multiple point measurements. In addition, the infrared gas analyzer requires artificial conditions with a high boundary layer conductance, which might deviate from the actual conditions that the plant experiences resulting in less representative measurements.
