## Additional file 3: Figure S2. for "Monitoring of drought stress and transpiration rate using proximal thermal and hyperspectral imaging in an indoor automated plant phenotyping platform"

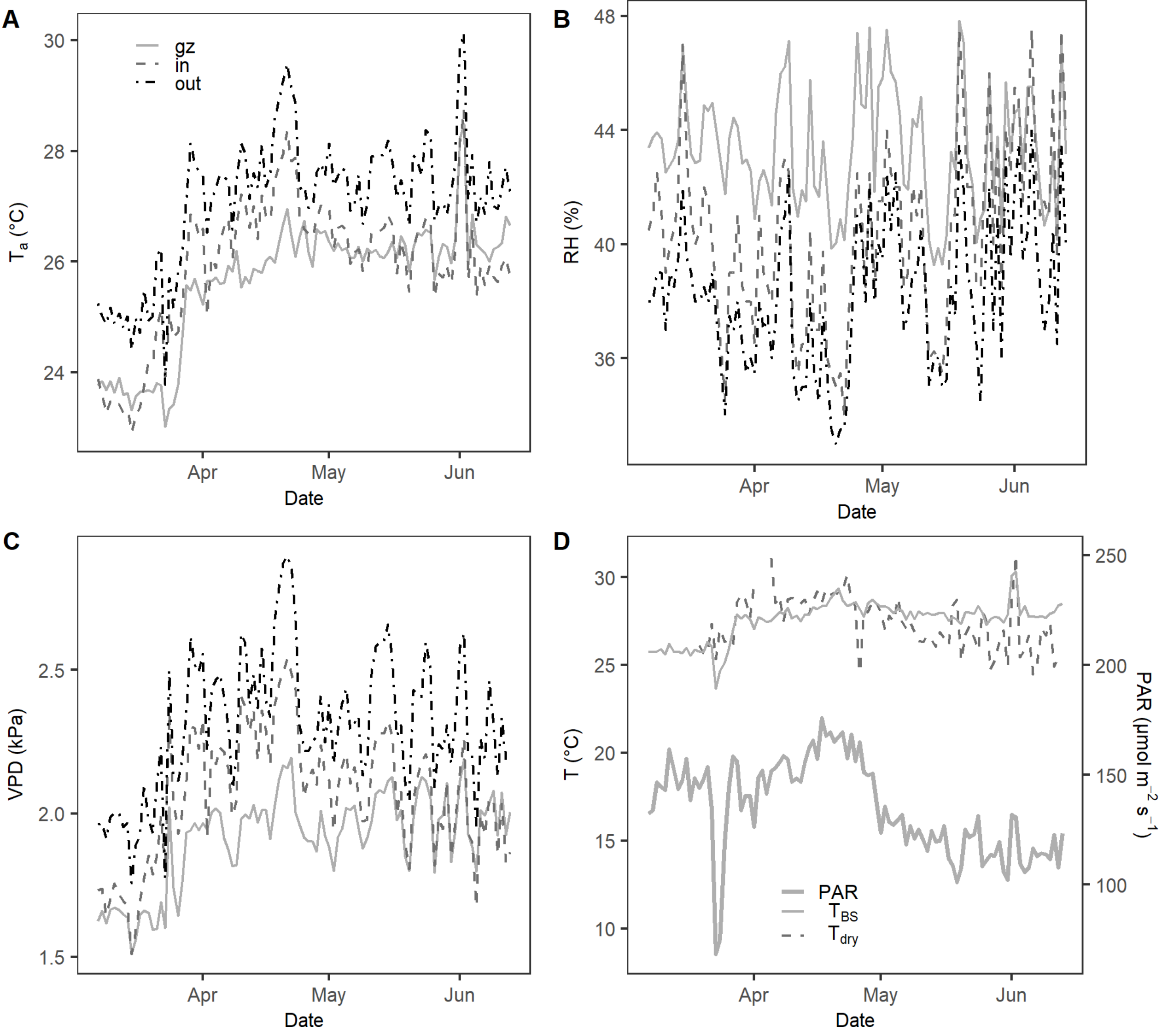


Environmental data. The daily mean air temperature (Ta, **A**), relative humidity (RH, **B**) and vapor pressure deficit (VPD, **C**) of the three monitoring positions (gz, in and out) are represented by a solid light gray line, dashed dark gray line and dotdashed black line, respectively. **D** shows the measurements of the black sphere temperature ($T_{BS}$), PAR monitored in the growth zone, and the dry reference temperature ($T_{dry}$) that was measured inside the imaging cabin. The daily mean of $T_{BS}$ and PAR are indicated by a thin and thick light gray line, respectively, while $T_{dry}$ is represented by a dashed dark gray line.
