## Additional file 4: Figure S3. for "Monitoring of drought stress and transpiration rate using proximal thermal and hyperspectral imaging in an indoor automated plant phenotyping platform"

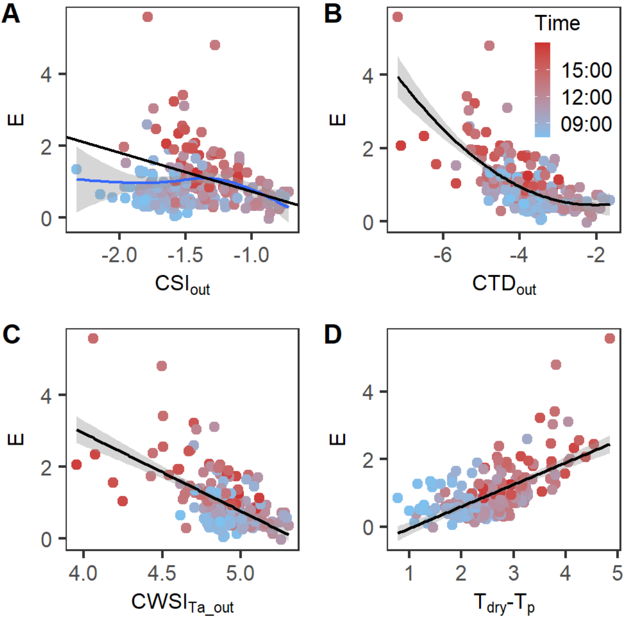


Relationship between thermal infrared indices and transpiration rate (E). Individual measurements are represented with colored dots showing the $VPD_{out}$ at the time of sampling. A blue-red gradient is used to visualize the time range. The linear or polynomial relationships between the indices are indicated with a black line, while non-linear (spline) relationships are represented by a blue line. The gray shading around the lines show the 95% confidence interval of the relationship. E (mmol $m^{-2}s^{-1}$) versus (**A**) $CSI_{out}$, (**B**) $CTD_{out}$, (**C**) $CWSI_{Ta\_out}$, and (**D**) $T_{dry}-T_p$ (°C).
